# When does parasitism maintain sex in the absence of Red Queen Dynamics?

**DOI:** 10.1101/2020.08.06.239632

**Authors:** Ben Ashby

## Abstract

Parasites can select for sexual reproduction in host populations, preventing replacement by faster growing asexual lineages. This is usually attributed to so-called “Red Queen Dynamics” (RQD), where antagonistic coevolution causes fluctuating selection in allele frequencies, which provides sex with an advantage over asex. However, parasitism may also maintain sex in the absence of RQD when sexual populations are more genetically diverse – and hence more resistant, on average – than clonal populations, allowing sex and asex to stably coexist. While the maintenance of sex due to RQD has been studied extensively, the conditions that allow sex and asex to stably coexist have yet to be explored in detail. In particular, we lack an understanding of how host demography and parasite epidemiology affect the maintenance of sex in the absence of RQD. Here, I use an eco-evolutionary model to show that both population density and the type and strength of virulence are important for maintaining sex, which can be understood in terms of their effects on disease prevalence and severity. In addition, I show that even in the absence of heterozygote advantage, asexual heterozygosity affects coexistence with sex due to variation in niche overlap. These results reveal which host and parasite characteristics are most important for the maintenance of sex in the absence of RQD, and provide empirically testable predictions for how demography and epidemiology mediate competition between sex and asex.

## Introduction

The question of why so many organisms reproduce sexually rather than asexually has been a long-standing puzzle in evolutionary biology (Maynard Smith, 1978). All else being equal, an asexual population will grow at twice the rate of a dioecious sexual population which pays a “two-fold cost” of producing males. The advantage of sex is generally attributed to the rapid and continual generation of diversity through the processes of recombination and segregation, whereas asexual lineages usually rely solely on mutations to generate novel variation (Hill & Robertson, 1966; Williams, 1975; Maynard Smith, 1978). While producing diverse offspring is not intrinsically beneficial, it may be advantageous in a variable environment where there is no fixed optimal phenotype. For example, if there exists heritable variation in resistance to parasites, then sex represents a bet-hedging strategy: producing offspring with variation in resistance allows the population to survive invasion by faster-growing but less diverse or less adaptable – and hence on average, more susceptible – asexual populations (Hamilton, 1980; Lively, 2010a).

Parasitism has been shown to select for sex both theoretically (reviewed in Lively, 2010) and empirically (Lively, 1987; King *et al*., 2009; Morran *et al*., 2011). Yet many studies assume that the advantages of sex are only realised when host and parasite populations exhibit negative frequency-dependent selection, as this can result in cyclical allele frequency dynamics and hence a continually varying environment with no fixed optimal phenotype. This cycling in allele frequencies due to antagonistic coevolution is often referred to as Red Queen Dynamics (RQD) in the literature, although this is in fact a broader term which also encompasses cycles that are not driven by negative frequency-dependent selection, and successive selective sweeps (Brockhurst *et al*., 2014). Almost all theoretical models for the evolutionary maintenance of sex by parasites depend on RQD, and as such there is an extensive literature on the causes and consequences of these dynamics (reviewed in Lively, 2010; see also recent work including Gokhale *et al*., 2013; Luijckx *et al*., 2013; Ashby & Gupta, 2014; Ashby & King, 2015; Ashby & Boots, 2017; MacPherson & Otto, 2018). However, as demonstrated by Lively (2010b), RQD are neither necessary nor sufficient for the maintenance of sex by parasites. Indeed, Lively (2010b) showed that it is possible for sexual and asexual hosts to coexist in the absence of RQD when the cost of producing males is offset by lower disease prevalence, on average. Thus, while an asexual lineage may initially have a population growth rate advantage, this is eventually curtailed by higher levels of infection, which allows the two populations to coexist.

Surprisingly, this mechanism for maintaining sex in the absence of RQD has yet to receive much theoretical attention. In particular, we currently do not know how the epidemiology of the parasite and the demography of the host impact on this mechanism, and hence its generality as an explanation for the maintenance of sex in the absence of RQD. There is good reason to suspect that a number of demographic and epidemiological characteristics of the populations will heavily influence selection for or against sex through effects on disease prevalence and severity. Moreover, in nature hosts and parasites vary widely in epidemiology and demography, and so it is necessary to explore variation in these processes in more detail to determine precisely when the costs of sex are offset by lower disease prevalence, and to what extent. Both the costs and benefits of sex may vary substantially between populations, with Maynard Smith’s “twofold cost of sex” representing a baseline assumption with all else being equal (for a detailed discussion of the costs of sex, see Lehtonen *et al*., 2012). The fitness benefits of sex related to parasitism will depend on factors such as the type and strength of virulence, the rate of recovery, the transmissibility of the parasite, the strength of resource competition, and the natural turnover rate of the population.

Most theoretical models of the maintenance of sex by parasites lack explicit population dynamics (see Table 2 in Ashby & King, 2015), and as such are unable to capture how interactions between demographic and epidemiological factors mediate selection for or against sex through changes in disease prevalence and severity. Here, I use an eco-evolutionary model with diploid hosts and haploid parasites to investigate the impact of demographic and epidemiological characteristics of the populations on the maintenance of sex in the absence of RQD. Specifically, I explore how all pairwise interactions between demographic (birth rate, natural death rate, and resource competition) and epidemiological (mortality virulence, sterility virulence, transmission rate, and recovery rate) factors affect the frequency of sex in the host population following the invasion of an asexual lineage. Through numerical analysis of the model, I show which characteristics of the populations tend to facilitate the maintenance of sex (and at higher frequencies). In addition, while there is no inherent heterozygote advantage in the underlying genetics of the model, I show that the frequency of sex depends on the heterozygosity of the asexual population, with sex maintained at a higher frequency when the invading asexual lineage is homozygous at all loci.

**Table 1.**
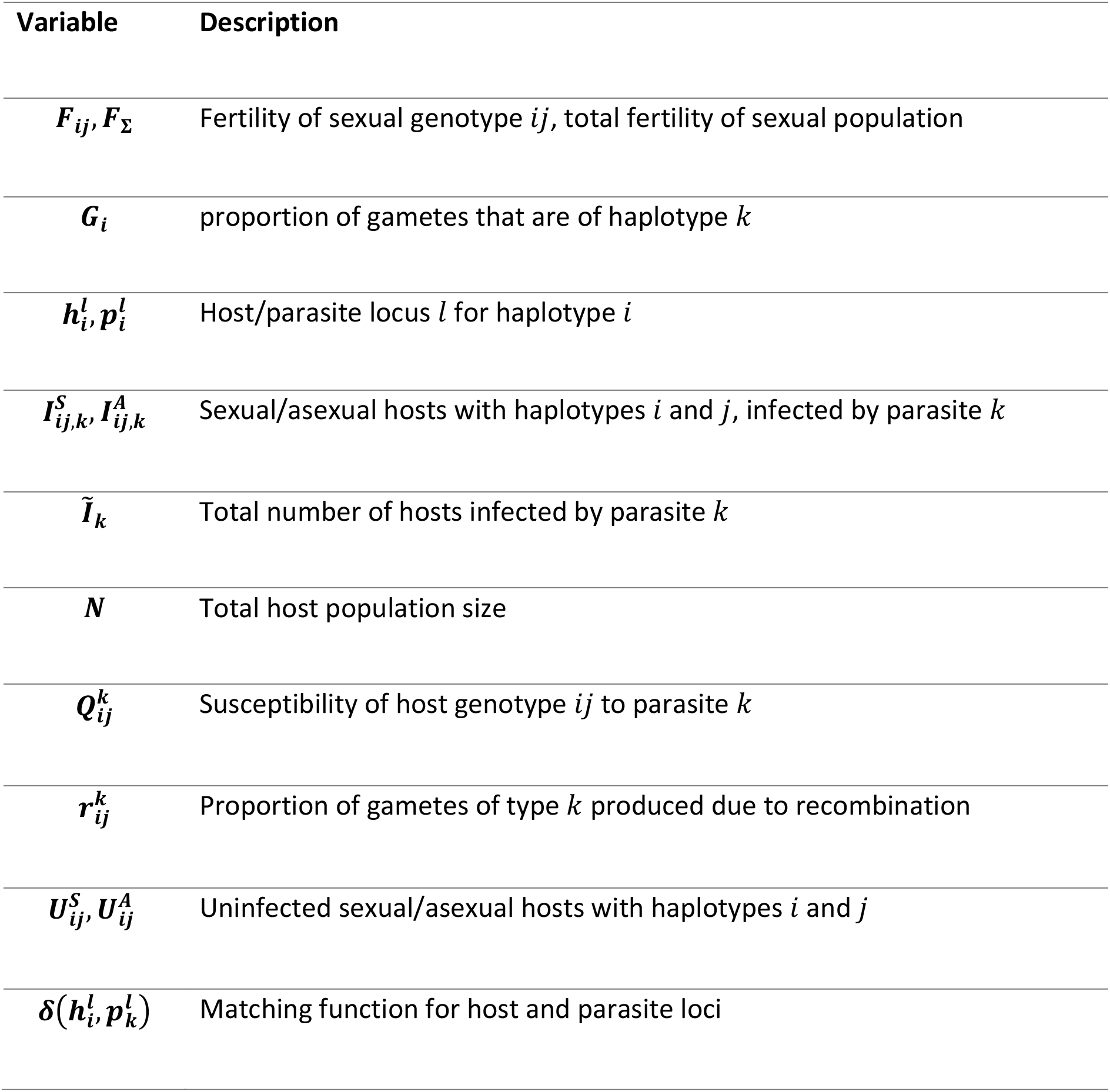
Description of model variables.

**Table 2.**
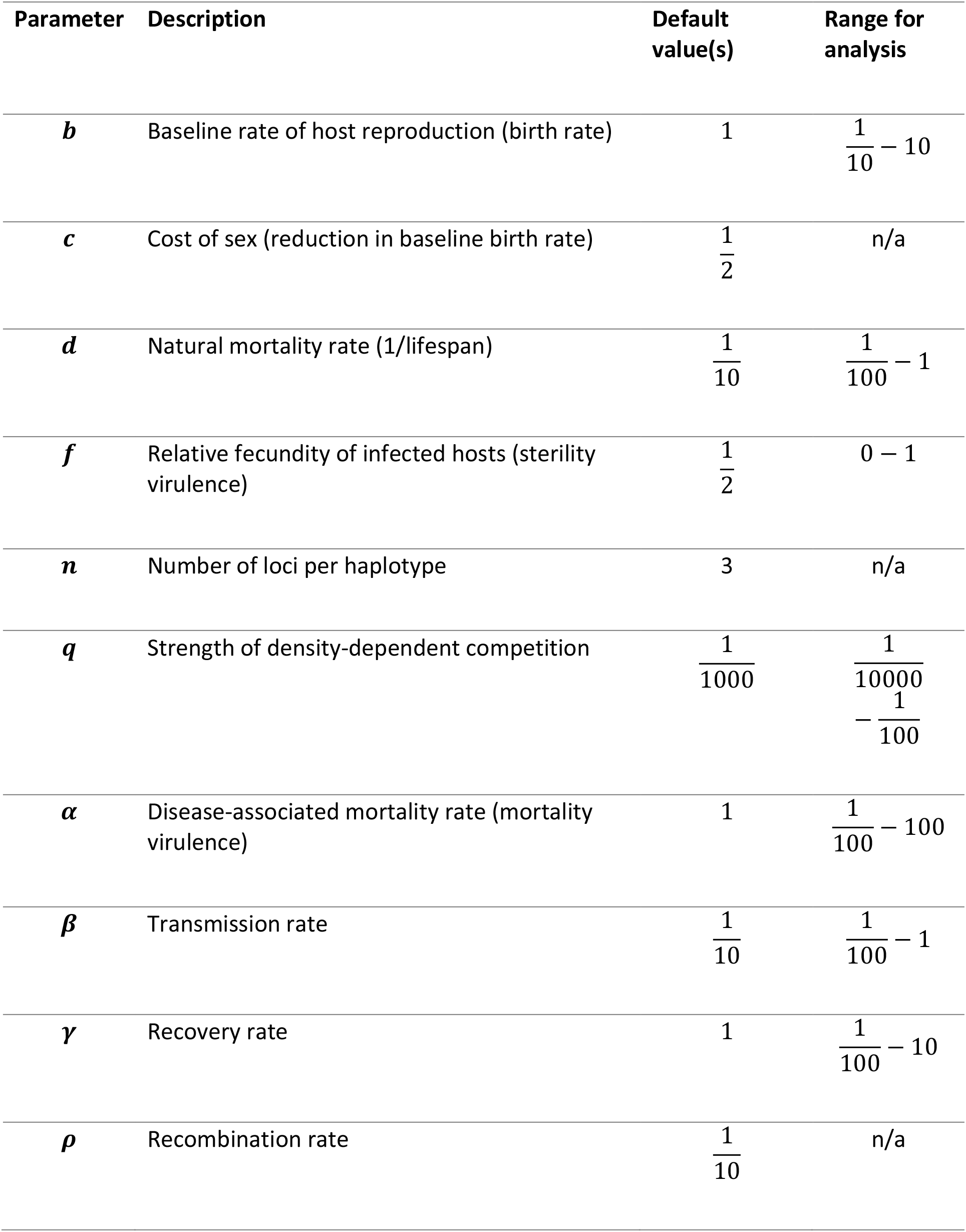
Model parameters, default values and ranges (where applicable).

| Parameter | Description | Default value(s) | Range for analysis |
| --- | --- | --- | --- |
| $b$ | Baseline rate of host reproduction (birth rate) | 1 | $\frac{1}{10} - 10$ |
| $c$ | Cost of sex (reduction in baseline birth rate) | $\frac{1}{2}$ | n/a |
| $d$ | Natural mortality rate (1/lifespan) | $\frac{1}{10}$ | $\frac{1}{100} - 1$ |
| $f$ | Relative fecundity of infected hosts (sterility virulence) | $\frac{1}{2}$ | 0 – 1 |
| $n$ | Number of loci per haplotype | 3 | n/a |
| $q$ | Strength of density-dependent competition | $\frac{1}{1000}$ | $\frac{1}{10000} - \frac{1}{100}$ |
| $\alpha$ | Disease-associated mortality rate (mortality virulence) | 1 | $\frac{1}{100} - 100$ |
| $\beta$ | Transmission rate | $\frac{1}{10}$ | $\frac{1}{100} - 1$ |
| $\gamma$ | Recovery rate | 1 | $\frac{1}{100} - 10$ |
| $\rho$ | Recombination rate | $\frac{1}{10}$ | n/a |

## Methods

I deterministically model the eco-evolutionary dynamics of diploid sexual and asexual hosts (superscripts *S* and *A*, respectively), and haploid asexual parasites. The populations are well-mixed and mating occurs randomly between sexual hosts. Transmission occurs horizontally from infected (*I*) to uninfected (*U*) hosts (there is no co-infection or super-infection) at a baseline pairwise transmission rate of *β*, with the susceptibility of the recipient, 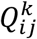, determined by the similarity of host (*ij*) and parasite (*k*) genotypes. Host genotypes are represented by binary strings of length 2*n* (i.e. for host type *ij* comprised of haplotypes *i* and 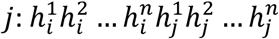, with 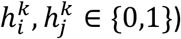 and parasite genotypes by binary strings of length *n* (i.e. for parasite type 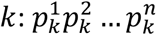, with 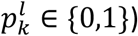. Note that heterozygotes *ij* and *ji* are phenotypically identical but are tracked separately for simplicity. The infection genetics follow a matching alleles framework (Frank, 1993), with host susceptibility equal to the average proportion of parasite loci which match the corresponding host loci on each host haplotype (Ashby & King, 2015):

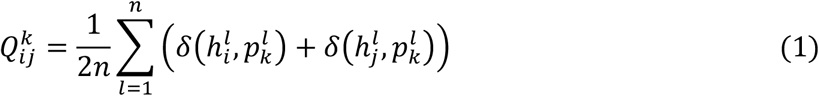

where 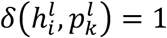 if 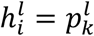. and 0 otherwise (hence 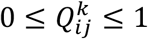). This means that heterozygotes are on average as susceptible to infection as homozygotes 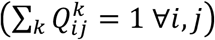. Hosts die naturally at rate *d* and from infection at rate *α* (mortality virulence), and may also suffer from reduced fertility due to infection, *f* (sterility virulence), with 0 ≤ *f* ≤ 1. Hosts recover from infection at rate *γ*.

Asexual hosts reproduce at a per-capita rate of *b*(1 − *qN*) where *b* is the baseline rate of reproduction (hereafter “birth rate”), *q* controls the strength of density-dependent competition, and *N* is the total population size of all hosts. Sexual hosts reproduce at a reduced baseline per-capita rate of *bc*(1 − *qN*), where 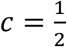 corresponds to a two-fold cost of sex. I assume that sexual hosts are always able to find a fertile mate, which means there are no additional costs of sex compared to asex due to unsuccessful mating. The fertility of sexual type *ij* is therefore 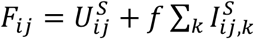 and the total fertility of the sexual population as 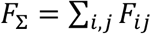. The proportion of gametes that are of haplotype *k* is 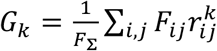, where 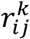 is the proportion of gametes of type *k* produced due to recombination, which occurs at a rate of *ρ* between neighbouring loci. The rate at which new sexual hosts of type *ij* are produced is therefore *bc*(1 − *qN*)*F*_Σ_*G_i_G_j_*.

The eco-evolutionary dynamics are described by the following set of ordinary differential equations:

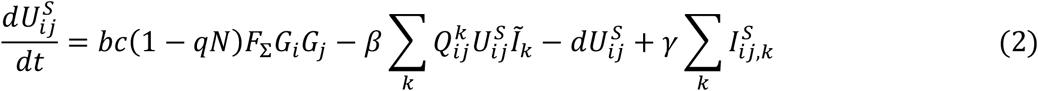

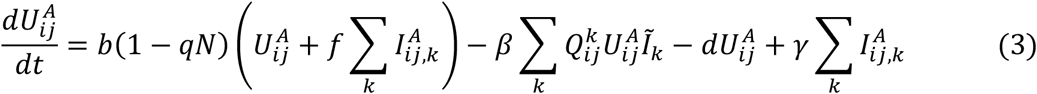

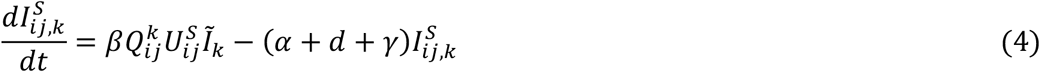

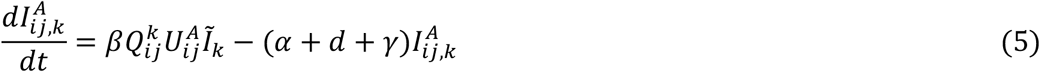

where 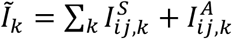 is the total number of hosts infected by parasite *k*.

To determine the effects of variation in epidemiological (*α, β, γ, f*) and demographic (*b, d, q*) parameters on the evolutionary maintenance of sex, I consider the invasion of a single asexual lineage into an established sexual population with all parasite genotypes initially present. The converse scenario – invasion of a sexual population from rare – would concern the evolutionary origins of sex rather than its maintenance, which is beyond the scope of the present study. To avoid unnecessary replication, I group asexual lineages with similar genotypes based on the sum of their binary strings, and consider one lineage from each group, scaling the results by the number of lineages within each group to represent the average over all asexual lineages. For example, if *n* = 2, each host has genotype of length 2*n* = 4, and there are five groups of asexual lineages to consider, with binary strings summing from zero to four: one asexual lineage in the first group (0000), four in the second (0001, 0010, 0100, 1000), six in the third (0011, 0101, 0110, 1001, 1010, 1100), four in the fourth (0111, 1011, 1101, 1110) and one in the fifth (1111). I allow the model to run for a total of *T_max_* = 10^4^ time units (time units are arbitrary, and longer durations do not qualitatively change the results), and measure the mean frequency of the sexual population and mean disease prevalence over the final 20% of each model run.

## RESULTS

### Epidemic thresholds

Suppose the population consists only of sexual hosts. In the absence of disease, the total population size will tend to 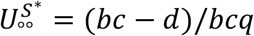 provided *bc > d* (if *bc < d* then the population will die out, so I assume that the disease-free sexual population is always viable). In this instance, selection is neutral among the various sexual genotypes, which means there are an infinite number of equilibria with 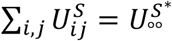. When the population is evenly distributed over all genotypes, any parasite is able to spread if the basic reproductive ratio, 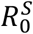 – the average number of secondary infections produced by a single infectious individual in an otherwise uninfected population – satisfies

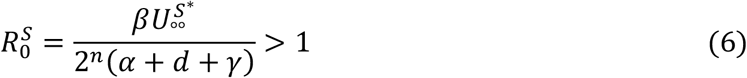

Similarly, in the absence of disease a population consisting of a single asexual lineage will tend to 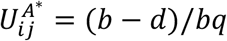 provided *b > d*, and parasite *k* is able to spread from rare when its basic reproductive ratio, 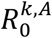, satisfies

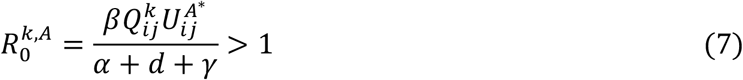

Clearly 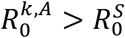 owing to the greater diversity of the sexual population, which limits transmission of the parasite. In general, parasite genotype *k* is able to spread in an entirely susceptible mixed host population when

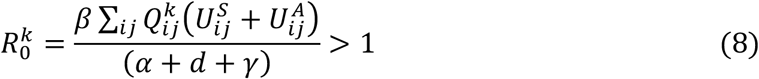

Note that by definition 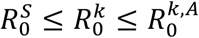. In the numerical analysis that follows, I initialise the sexual host population close to its endemic equilibrium (omitted here as it is too unwieldy) when 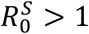, and close to the disease-free equilibrium (with all parasites rare) when 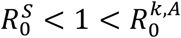.

### Asexual heterozygosity determines the frequency of sex

I now consider the invasion of a single, rare asexual lineage into an established sexual population. Intuitively, there is always at least one asexual lineage that can invade a viable sexual population when there is a cost of sex (*c* < 1), regardless of the genetic distribution of the population. This is because sexual and asexual hosts are equivalent up to reproduction, and since at least one sexual genotype must have non-zero growth rate for the population to be viable, there is a corresponding asexual genotype which has a positive growth rate when rare (since *c* < 1). This means that the sexual population cannot drive the asexual population extinct deterministically (but could do so stochastically). Instead, the asexual lineage will either deterministically exclude the sexual population, or the two populations will coexist at a stable equilibrium.

Crucially, the choice of asexual lineage can have a significant impact on the outcome, with the sexual population always at a higher frequency when the asexual lineage is homozygous at all loci (i.e. it has a repeated haplotype; Fig. 1). In some cases, the sexual population will stably coexist with an asexual lineage that is fully homozygous but is driven extinct if the asexual lineage is heterozygous at any loci (Fig. 1a). Yet this is not due to heterozygote advantage, as all host genotypes are equally susceptible, on average. The difference in outcomes occurs because individuals with repeated haplotypes, which are fully susceptible to a single parasite, have a narrower niche than individuals with non-repeated haplotypes, which are half as susceptible to two parasites. The extent to which the sexual and asexual populations compete depends on the degree of niche overlap between them: all haplotypes are initially represented in the sexual population, but only one (repeated haplotype) or two (distinct haplotypes) occur in the asexual population. Thus, when there is a single repeated haplotype in the asexual population, the degree of niche overlap with the sexual population, and hence the strength of competition, is lower, allowing sex to be maintained at a higher frequency. For example, in Fig. 1a there is a single locus, so there are three possible host genotypes: 0/0, 0/1, and 1/1. If the invading asexual genotype is homozygous (0/0 or 1/1), then it coexists with the sexual population. In the sexual population, the heterozygote (0/1; partial niche overlap) and the non-matching homozygote (1/1 or 0/0; no niche overlap with respect to parasitism) are maintained at moderate frequencies, whereas the matching homozygote (0/0 or 1/1; full niche overlap) is only maintained at a low frequency due to segregation. If, however, the asexual genotype is heterozygous (0/1) then it has partial or full niche overlap with all sexual hosts, which are driven extinct as a result. As the number of loci increases, the asexual genotype always overlaps with a smaller and smaller proportion of the sexual population, and so the difference in outcomes between homozygous and heterozygous asexual lineages reduces (Fig. 1b-c).

**Figure 1.**
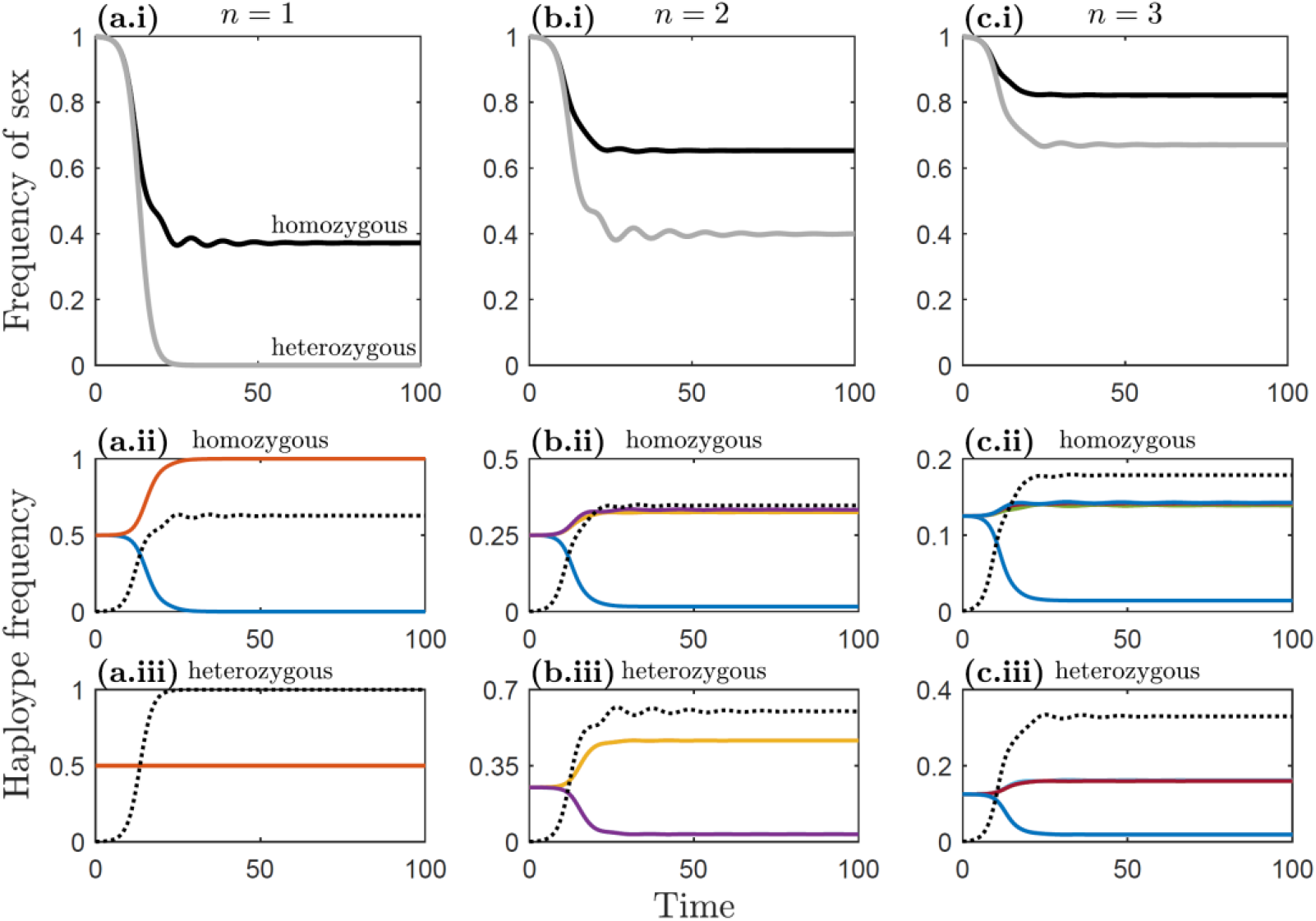
Example model dynamics for different numbers of loci per haplotype, *n*: (a) *n* = 1; (b) *n* = 2; (c) *n* = 3. Top row (a.i-c.i): Sex persists at a higher frequency when the invading asexual host is fully homozygous (black) rather than heterozygous (grey), and as the number of loci increases. Middle row (a.ii-c.ii): Haplotype frequencies for sexual (solid, colour) and asexual (dotted, black) haplotypes when the invading asexual host is homozygous at all loci. Bottom row (a.iii-c.iii): As middle row, but when the invading asexual host is heterozygous at all loci. In all cases, the long-term dynamics tend to a stable equilibrium, with any cycles (Red Queen Dynamics; RQD) being transient. Fixed parameters as described in Table 2, except *γ* = 0.1 to demonstrate transient RQD.

### Demography and epidemiology affect the maintenance of sex in the absence of Red Queen Dynamics

I now consider how variation in host demography (birth rate, *b*; death rate, *d*; competition, *q*) and parasite epidemiology (mortality virulence, *α*; sterility virulence, *f*; transmission rate, *β*; recovery rate, *γ*) affect the frequency of sexual reproduction in the host population by exploring all pairwise combinations of parameters. I solve the model numerically as it is analytically intractable in all but the simplest of cases, with parameters ranges as described in Table 2. The numerical results are presented in Fig. 2-7 and are summarised in Table 3.

**Table 3.** Summary of model conditions which are generally more favourable to sex.

| Parameter | Conditions generally more favourable to sex | Notable exceptions |
| --- | --- | --- |
| Birth rate, $b$ | low/intermediate birth rates | small $\beta$; large $q$, $\alpha$ or $\gamma$ |
| Natural mortality rate, $d$ | low natural mortality rates (longer lifespans) | small $q$ or $\gamma$; large $\beta$ |
| Competition, $q$ | weak/moderate competition between hosts | small $\gamma$; large $b$ or $\beta$ |
| Mortality virulence, $\alpha$ | intermediate mortality virulence | - |
| Sterility virulence, $f$ | stronger sterility virulence ($f \ll 1$) | - |
| Transmission rate, $\beta$ | high/intermediate transmission rates | - |
| Recovery rate, $\gamma$ | low/intermediate recovery rates | large $f$ |

**Figure 2.**
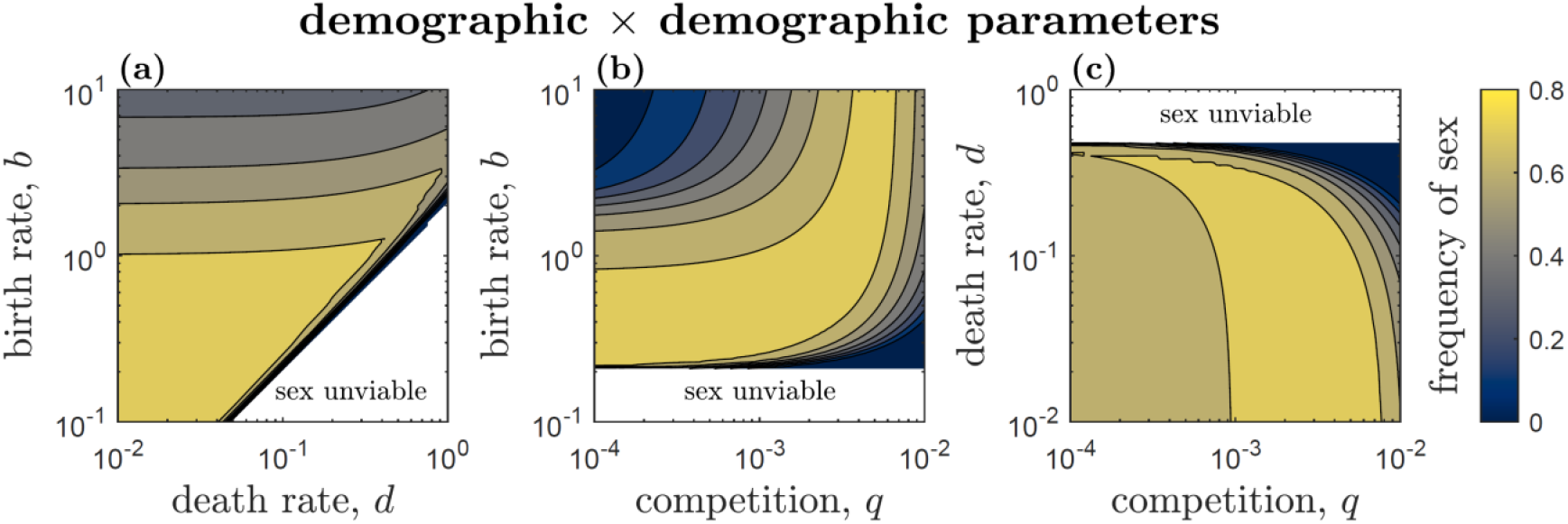
Effects of interactions between demographic parameters (birth rate, *b*; death rate, *d*; competition, *q*) on the frequency of sex. Panels show the frequency of sex at equilibrium after the invasion of a single asexual lineage, averaged over all possible asexual lineages. Contours and shading represent 0.1 increments in the frequency of sex (as a proportion of the population), with the lightest region corresponding to parameters which result in the frequency of sex being between 0.7 and 0.8 (i.e. 70-80%), and the darkest between 0 and 0.1 (i.e. 0-10%). Sex is unviable in the white region. Fixed parameters as described in Table 2.

Excluding trivial outcomes (e.g. sexual hosts are unviable), an invading asexual lineage either excludes the sexual population, or they coexist at a stable equilibrium. I found no instances where the populations exhibited persistent Red Queen Dynamics (fluctuating selection). Instead, any cycling in haplotype frequencies was damped, and hence transient (Fig. 1). Unlike most previous models, when sex is maintained it is not due to Red Queen Dynamics but is instead due to the costs of sex being offset by having a lower risk of infection, on average (Lively, 2010b). Numerical analysis of the model reveals that this trade-off is sufficient to maintain sex across a wide range of demographic and epidemiological parameters.

#### Demographic × demographic parameters

All three demographic parameters in the model have a considerable impact on competition between sexual and asexual hosts, with non-linear pairwise interactions between them (Fig. 2). The frequency of sex peaks when population turnover is slow (i.e. low birth (*b*) and natural death (*d*) rates), with sex faring worse as the birth rate increases or at relatively high natural death rates, just before the sexual population becomes unviable (Fig. 2a). There is a non-linear interaction between the birth rate and the strength of resource competition (*q*), with sex maintained at a high frequency when birth rates are low and resource competition is weak, or when birth rates are higher and resource competition is strong (Fig. 2b). This pattern occurs due to the effects of demography on the density of the host population, which modulates the risk of infection and hence the benefits of sex (Fig. 3b). When the population is too large (high birth rates and weak resource competition), most individuals are infected regardless of whether they are sexual or asexual, thus negating the benefits of sex. When the population is too small (low birth rates and strong resource competition), few individuals are infected and so the risk of infection is negligible, which also selects against sex. When the demographic parameters lead to populations of intermediate size, the risk of infection is high enough to allow sex to be maintained, but not so high that most sexual hosts are also infected. Finally, the frequency of sex generally decreases as the natural mortality rate increases, except when competition is relatively weak, where it peaks at intermediate values of *d* (Fig. 2c). Regions where sex is at high frequency correspond with low to moderate levels of disease in the population (^~^10-30%; Fig. 3), and therefore most members of the sexual population are uninfected.

**Figure 3.**
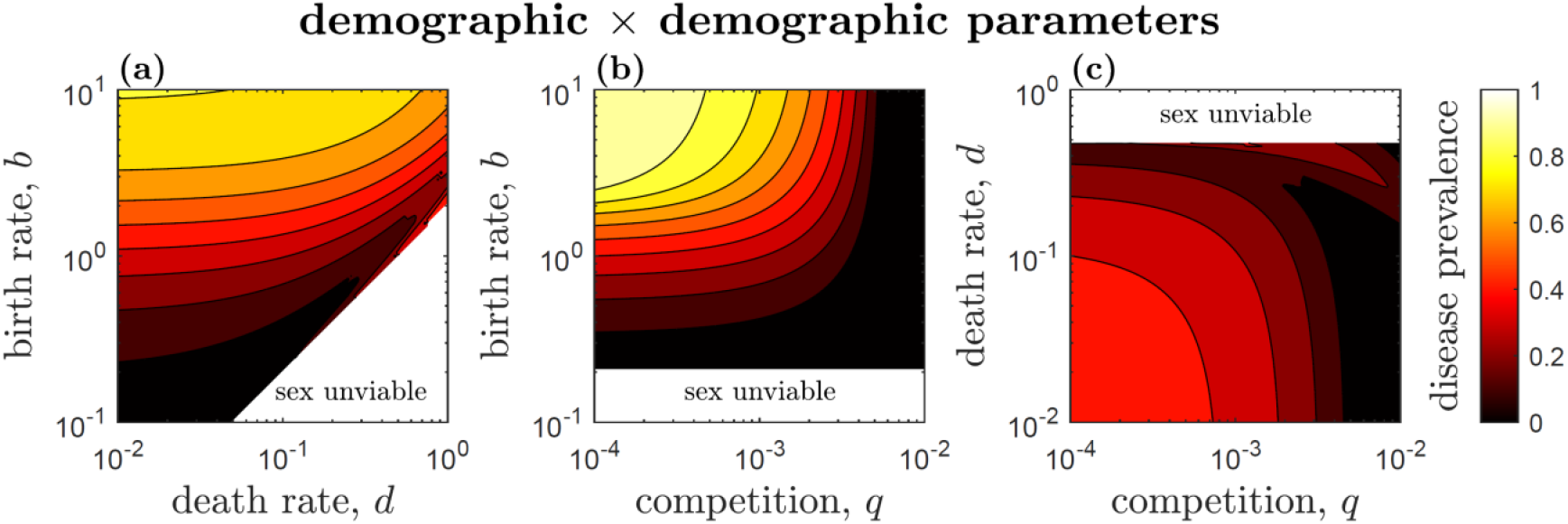
Effects of interactions between demographic parameters (birth rate, *b*; death rate, *d*; competition, *q*) on disease prevalence. Panels show the equilibrium disease prevalence after the invasion of a single asexual lineage, averaged over all possible asexual lineages. Contours and shading represent 0.1 increments in disease prevalence (as a proportion of the total population), with the lightest region corresponding to parameters which result in disease prevalence between 0.9 and 1 (i.e. 90-100%), and the darkest between 0 and 0.1 (i.e. 0-10%). Sex is unviable in the white region. Fixed parameters as described in Table 2.

#### Epidemiological × epidemiological parameters

All pairwise interactions between epidemiological parameters influence the frequency of sex, but to varying extents (Fig. 4). Most notably, mortality virulence (*α*) strongly interacts with all epidemiological parameters, but with the frequency of sex always peaking at intermediate mortality virulence (Fig. 4a-c). High mortality virulence means that the cost of infection is larger and hence there is a greater advantage to reproducing sexually. However, mortality virulence also lowers overall disease prevalence in the population by shortening the infectious period (Fig. 5a-c), and so the risk of being infected reduces as mortality virulence increases. Thus, the frequency of sex peaks for intermediate mortality virulence. In contrast, while sterility virulence (*f*) interacts strongly with mortality virulence (Fig. 4a), it has a much weaker impact on the frequency of sex as the transmission or recovery rate vary (Fig. 4d-e). Lastly, as the recovery rate increases a larger transmission rate is required to maintain sex at high frequency, which is due to their opposing effects on disease prevalence (Fig. 4f, 5f).

**Figure 4.**
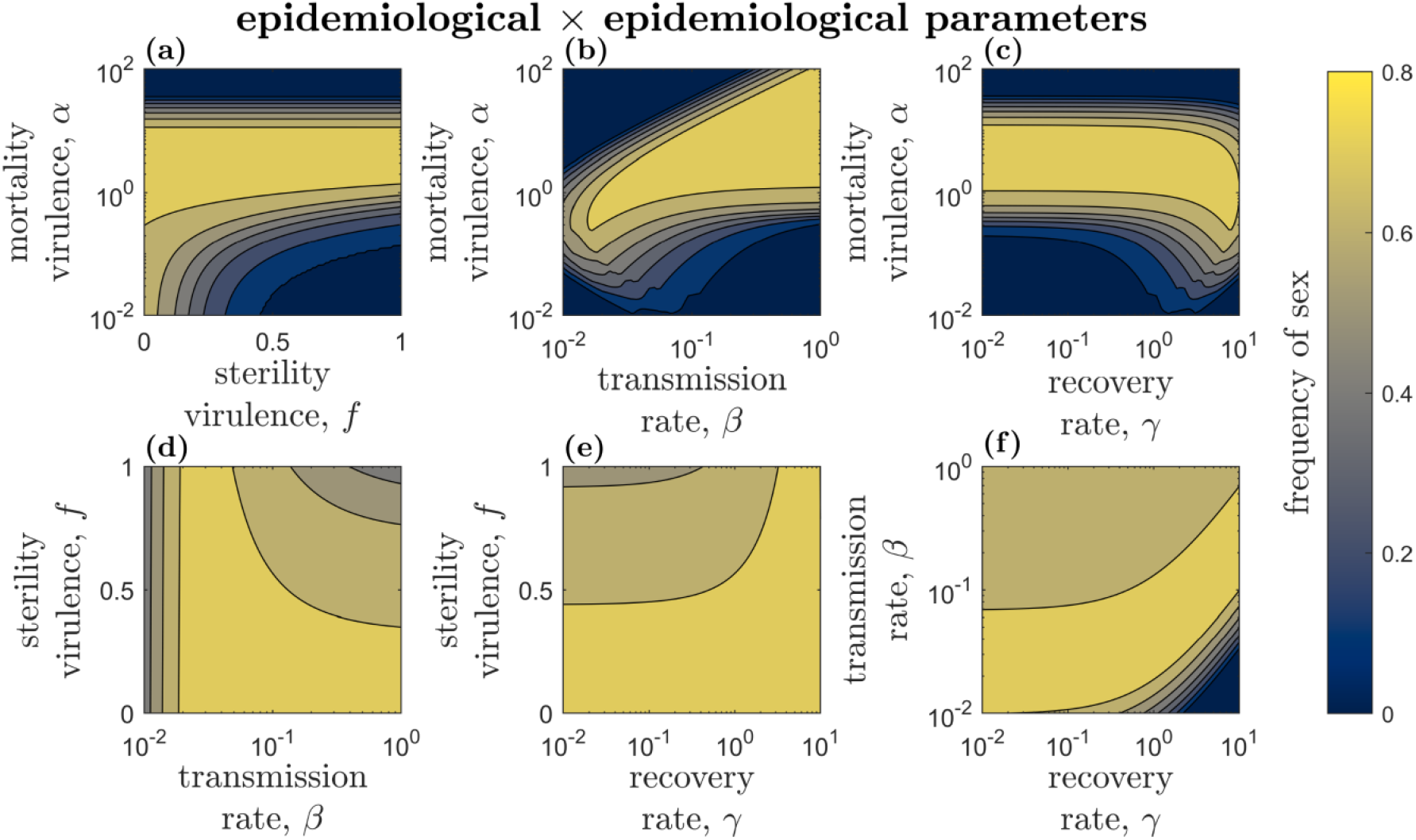
Effects of interactions between epidemiological parameters (mortality virulence, *α*; sterility virulence, *f*; transmission rate, *β*; recovery rate, *γ*) on the frequency of sex. Contours and shading as described in Fig. 2. Fixed parameters as described in Table 2.

**Figure 5.**
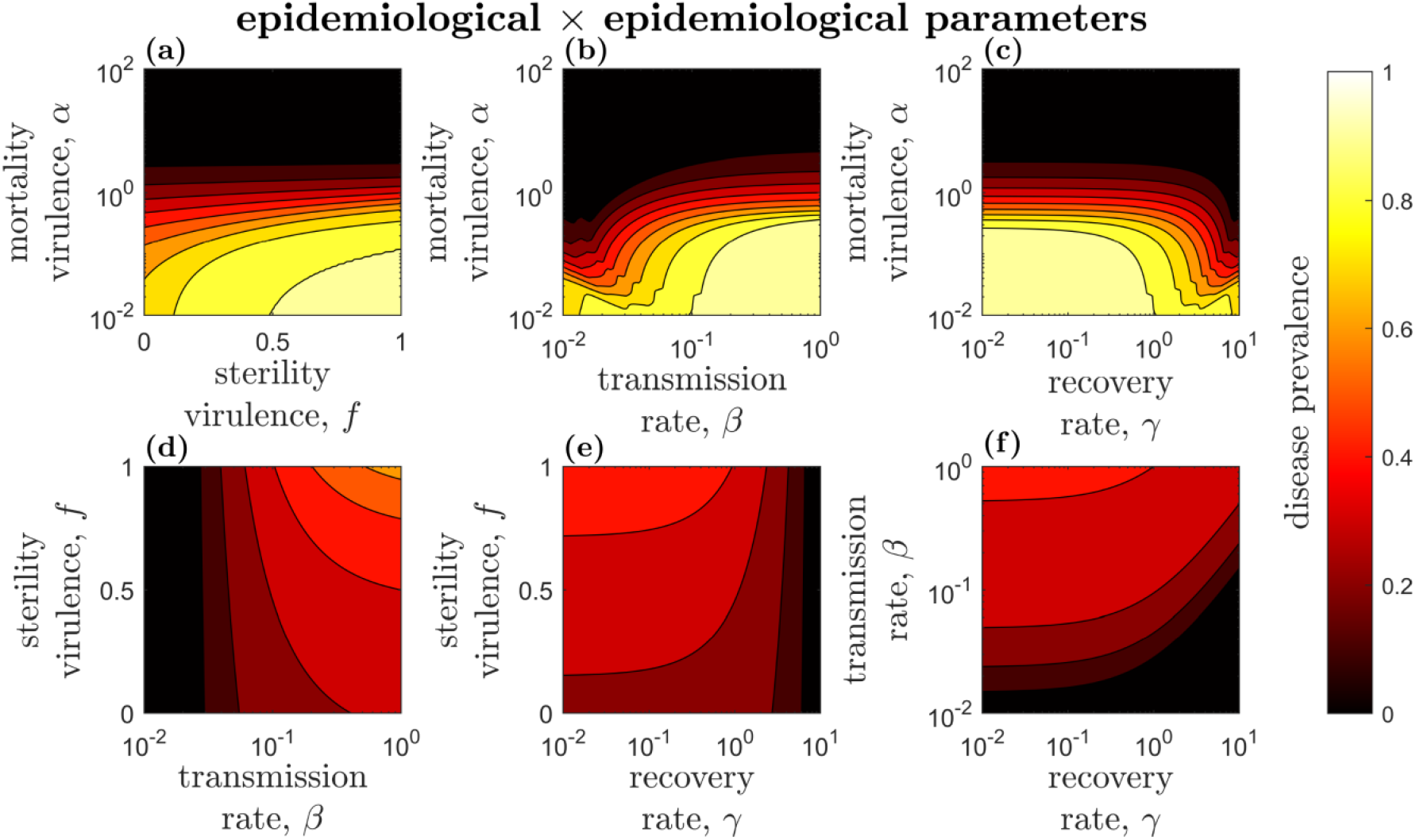
Effects of interactions between epidemiological parameters (mortality virulence, *α*; sterility virulence, *f*; transmission rate, *β*; recovery rate, *γ*) on disease prevalence. Contours and shading as described in Fig. 3. Fixed parameters as described in Table 2.

#### Demographic × epidemiological parameters

As there are 12 combinations consisting of one demographic and one epidemiological parameter, here I summarise broad patterns in the results (Fig. 6-7). Overall, the natural death rate and strength of resource competition mostly have similar interactions with the epidemiological parameters (Fig. 6b.i-c.iv), as they both reduce the host population size. Similarly, interactions between the demographic parameters and transmission and recovery are broadly similar, albeit reflected in the vertical axis due to their opposing effects on overall disease prevalence (Fig. 6a.iii-c.iv, 7a.iii-c.iv). Across the board, sex generally peaks when disease prevalence is roughly 10-30% (Fig. 7). In addition to these broad patterns, there are some notable pairwise interactions. For example, the birth rate strongly interacts with all the epidemiological parameters, and is the only parameter to have a substantial effect on the frequency of sex as sterility virulence varies (Fig. 6a.ii). This is likely because *f* always has *b* as a coefficient and so is directly modulated by *b* in the birth term of the model. The only other demographic parameter to have a noticeable (although much weaker) impact on the frequency of sex as sterility virulence varies is resource competition (Fig. 6c.ii), which also appears as a coefficient of some of the terms containing *f*.

**Figure 6.**
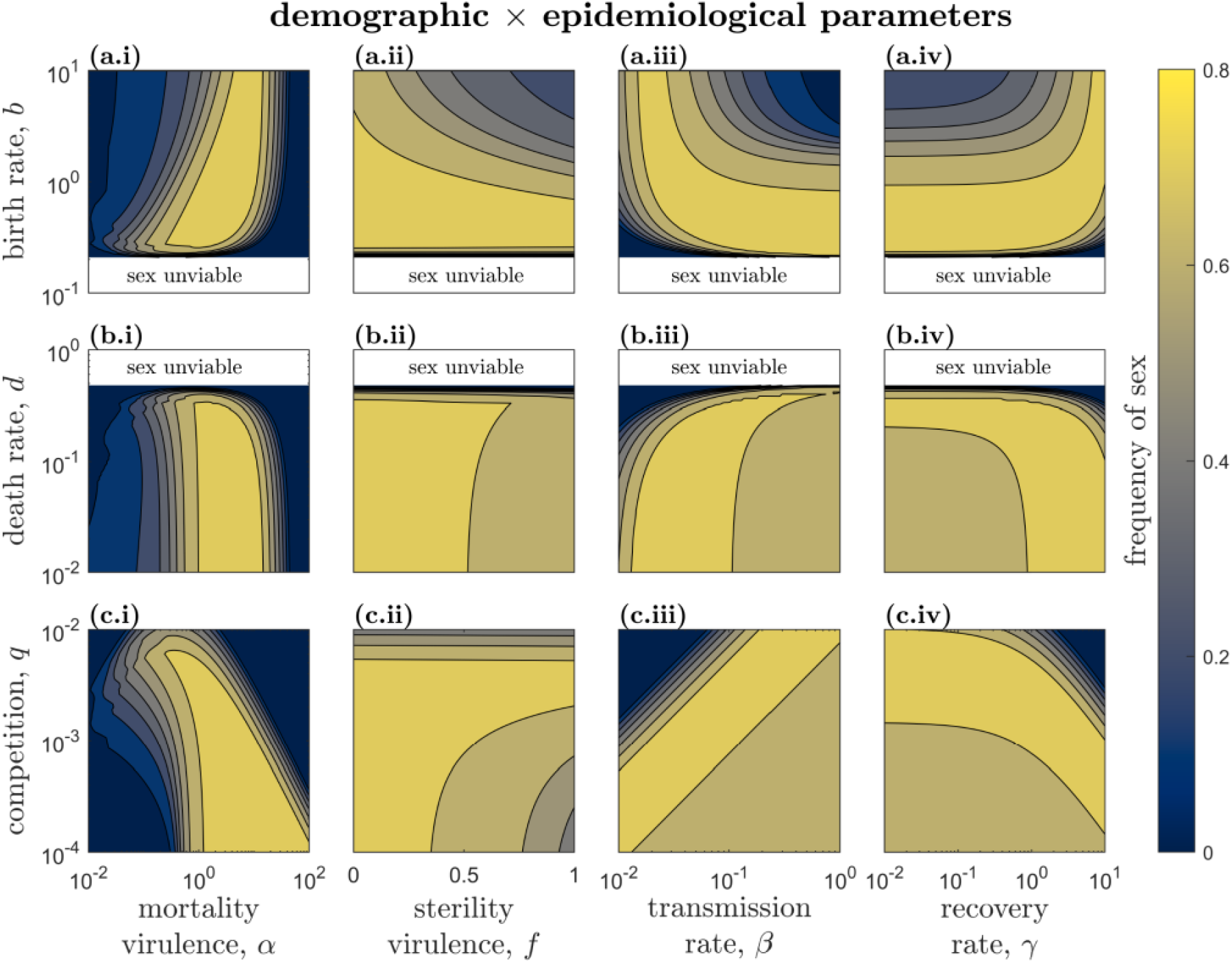
Effects of interactions between demographic (birth rate, *b*; death rate, *d*; competition, *q*) and epidemiological (mortality virulence, *α*; sterility virulence, *f*; transmission rate, *β*; recovery rate, *γ*) parameters on the frequency of sex. Sexual hosts are unviable in the white region. Contours and shading as described in Fig. 2. Fixed parameters as described in Table 2.

**Figure 7.**
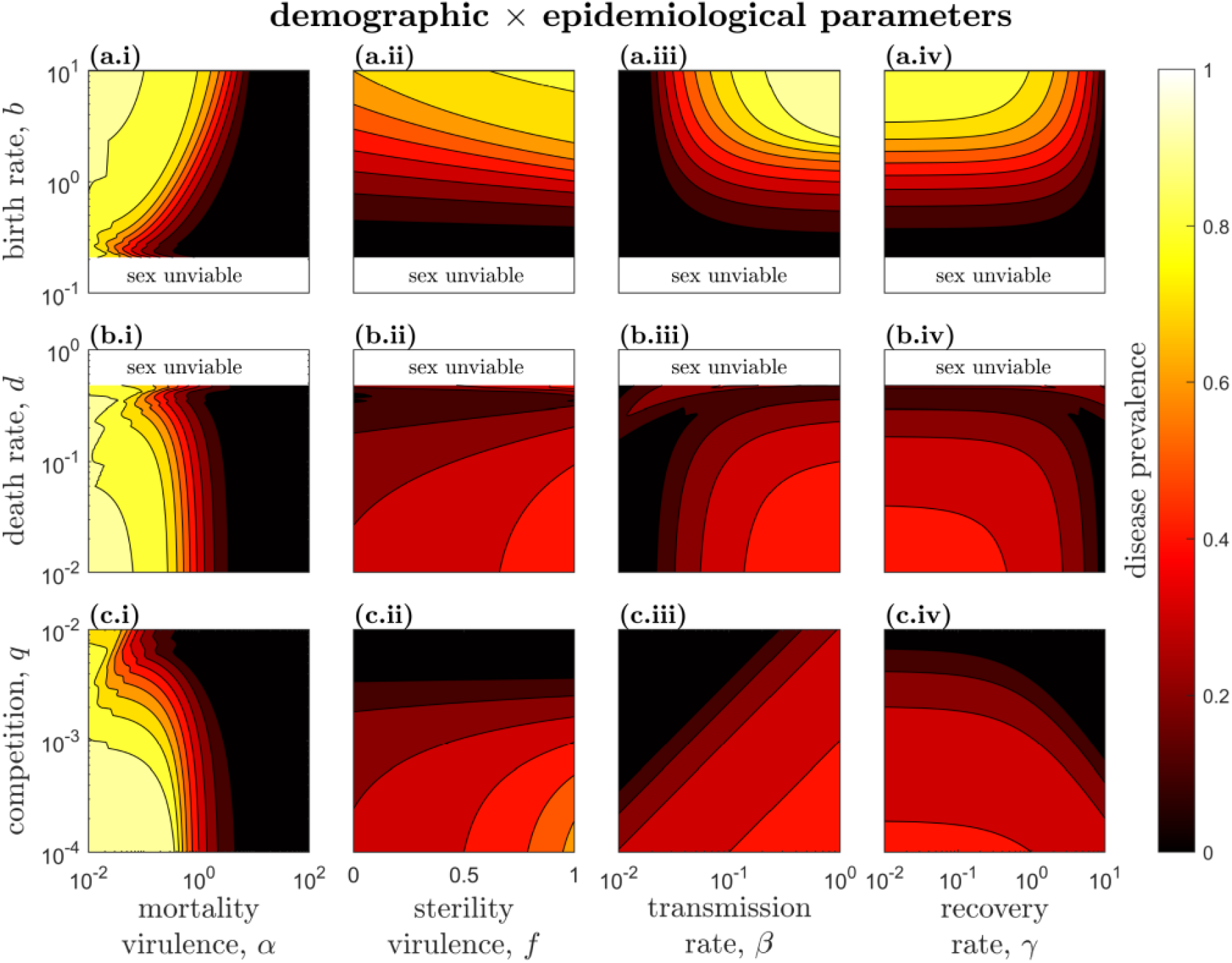
Effects of interactions between demographic (birth rate, *b*; death rate, *d*; competition, *q*) and epidemiological (mortality virulence, *α*; sterility virulence, *f*; transmission rate, *β*; recovery rate, *γ*) parameters on disease prevalence. Sexual hosts are unviable in the white region. Contours and shading as described in Fig. 3. Fixed parameters as described in Table 2.

## Discussion

Using a deterministic eco-evolutionary model of hosts and parasites, I have explored how interactions between demographic and epidemiological processes affect competition between sexual and asexual hosts. Unlike most previous models that rely on Red Queen Dynamics (RQD) – that is, co-evolutionary cycling of allele frequencies – to maintain sex (reviewed in Lively, 2010), here, sex and asex coexist at a stable equilibrium when the cost of producing males is offset by lower disease prevalence in the sexual population. The existence of this stable coexistence equilibrium, where the costs and benefits of sex are balanced, was first demonstrated by Lively, (2010b), but the extent to which host demography and parasite epidemiology combine to affect the maintenance of sex in the absence of RQD has not previously been explored.

Demographic factors such as the birth rate and strength of resource competition modulate disease prevalence – and hence selection for or against sex – through changes in population size. When the population size is too low, disease is rare and so selection favours asex because the risk of infection is low. When the population size is too high, disease is so common that most hosts are infected regardless of reproductive strategy. Thus, it is only at intermediate population sizes where disease is sufficiently prevalent that sex confers an advantage in terms of resistance, but not so prevalent that resistance becomes futile. This balance between disease prevalence and selection for parasite defence is common in the eco-evolutionary literature, and has previously been identified for the maintenance of sex (Ashby & King, 2015), as well as many other traits including recovery (van Baalen, 1998), mutualism (Rafaluk-mohr *et al*., 2018), and sociality (Bonds *et al*., 2005). The relationship between disease prevalence and the maintenance of sex once again highlights the importance of including density-dependent ecological/epidemiological dynamics in evolutionary models, rather than assuming a constant risk of infection as in classical population genetics frameworks, which often leads to qualitatively different results (Ashby *et al*., 2019).

The effects of parasite epidemiology on the frequency of sex can also be largely understood in terms of their impacts on disease prevalence, but also in terms of disease severity. These effects are most apparent when comparing the two modes of virulence – mortality and sterility. Sex peaks at intermediate mortality virulence due to a balance between disease prevalence, which decreases with greater virulence (shorter infectious period), and the cost of infection, which increases with virulence. Sterility virulence, on the other hand, does not alter the infectious period and so has a comparatively weak effect on disease prevalence, but does increase the cost of being infected. Sex therefore increases with sterility virulence (i.e. as *f* decreases), although the effect is generally weak compared to variation in most other parameters. Sterility virulence has a rather weak effect because the cost of infection appears to be dominated by mortality virulence. When mortality virulence is low, variation in sterility virulence has a much stronger effect on the frequency of sex (Fig. 4a). This suggests that when parasites cause appreciable levels of both mortality and sterility, the former should play the greater role in mediating competition between sexual and asexual hosts. However, there are likely to be two notable exceptions to this rule. First, if any reduction in host fecundity due to infection is permanent, then the costs of sterility virulence will be much higher, potentially increasing the advantages of sex over asex. Second, if hosts are unable to find a fertile mate, then sexual individuals will pay an additional cost of infection that asexuals do not incur. To see why, consider sexual and asexual populations each with average fertility *F*, where 0 < *F* < 1. If each sexual host mates randomly once per unit time, then the number of sexual offspring produced is proportional to *F*^2^ < *F*. In reality, individuals may mate multiple times and choose their mates based on quality or condition, including infection status or perceived levels of resistance (Hamilton & Zuk, 1982; Loehle, 1997; Kokko *et al*., 2002; Howard & Lively, 2003; Ashby & Boots, 2015; Ashby, 2020). For the sake of simplicity, the model examined herein assumed that there was no additional cost of sex due to sterility virulence, but this cost has previously been explored in the context of RQD, with sterility virulence increasing RQD but imposing an additional cost on sex (Ashby & Gupta, 2014). In summary, while the effects of sterility virulence in the present study are generally rather weak, there are certain circumstances where it may play a major role: specifically, when sterility is permanent or when it imposes an additional cost on sex.

The model also revealed how interactions between host demography and parasite epidemiology affect the frequency of sex. Again, the effects of these interactions on competition between sex and asex (Fig. 6) can largely be understood in terms of changes in disease prevalence (Fig. 7). Hence, certain epidemiological parameters, such as transmission and recovery rates, had broadly similar interactions with demographic parameters (albeit reversed in this instance due to their opposing effects on disease prevalence). Overall, the analysis indicated that the birth rate and mortality virulence are likely to be particularly important for modulating the frequency of sex in the population, as they interacted strongly with most parameters.

The present study has primarily focused on interactions between demographic and epidemiological processes, but other factors may also play an important role in the maintenance of sex in the absence of RQD. For instance, while there is no inherent heterozygote advantage in the model, the outcome of competition between sexual and asexual populations depends on the choice of asexual lineage. This can be understood in terms of niche overlap with respect to susceptibility, as the asexual population suppresses sexual genotypes with which it shares one or more haplotypes. Asexual lineages with a repeated haplotype (i.e. homozygous at all loci) are only susceptible to one parasite and so have a narrower niche overlap with the sexual population, whereas lineages with nonrepeated haplotypes are half as susceptible to two parasites and so have a broader niche overlap. Competition is therefore stronger in the latter case, which leads to a lower frequency of sex overall. The relative diversity of the sexual and asexual populations is also known to be crucial for the maintenance of sex (Ashby & King, 2015). In the present study, the asexual population only consisted of a single lineage, with relative diversity controlled by the number of loci. The number of loci was fixed in the main analysis because it did not qualitatively change the effects of demographic and epidemiological processes on the frequency of sex, which were the primary focus of the study. In real populations, the relative diversity of sexual and asexual hosts will also be controlled by rates of migration, mutation, recombination, and local extinction. Since the model was deterministic, these factors either had a negligible effect (recombination) or were omitted for simplicity (migration, mutation, extinction). In a stochastic model genotypes may be lost and reintroduced through these processes, and so they will affect the relative diversity of the populations (Gokhale *et al*., 2013; Ashby & King, 2015). Future work should explore the extent to which increasing the diversity of the asexual population reduces the frequency of sex. Similarly, while the present study focused on the evolutionary maintenance of sex by considering asexual invasion into an established sexual population (equivalent to rare migration or mutation), future work should also consider the converse scenario – which is more pertinent to the origin of sex – as the sexual population may have relatively low diversity when rare.

Another important factor in the maintenance of sex is the nature of the underlying infection genetics. That is, the degree to which each host genotype is susceptible to each parasite genotype. To date, the role of infection genetics in the maintenance of sex has almost exclusively been considered in the context of RQD. “Matching Alleles” frameworks (Frank, 1993; Luijckx *et al*., 2013) tend to produce large amplitude, high frequency cycles driven by negative frequency-dependent selection among specialists, whereas “Gene-for-Gene” frameworks (Parker, 1994; Sasaki, 2000) may produce slower elliptical cycles due to fitness costs associated with varying degrees of generalism. Although traditionally these paradigms have been considered as two ends of a continuum (Agrawal & Lively, 2002), Matching Alleles can also be considered as a subset of the Gene-for-Gene framework (Ashby & Boots, 2017). Nevertheless, few models in the literature assume Gene-for-Gene infection genetics, likely because early studies suggested the nature of the resulting RQD was generally not favourable to sex (Parker, 1994; Otto & Nuismer, 2004). Yet since sex can be readily maintained in the absence of RQD, the Gene-for-Gene framework deserves further attention in future theoretical models of sex versus asex.

The results have two implications for experimental work. First, the model makes a number of qualitative predictions for how variation in host or parasite life-history traits will affect selection for sex. It is important to note that the modelling predictions are predicated on eco-evolutionary feedbacks, and so any experiments must allow for disease prevalence and hence the risk of infection to vary naturally, rather than controlling exposure externally. In principle, one could readily test the prediction that sex peaks at intermediate mortality virulence by challenging mixtures of sexual and asexual populations with parasites that vary in their effects on host mortality. To the best of my knowledge, there have yet to be any experimental tests of how demographic or epidemiological traits affect the maintenance of sex by parasites. Indeed, opportunities for direct empirical tests of the maintenance of sex by parasites are limited by the need for comparable host populations that reproduce sexually and asexually. Many fungi and protists switch between sex and asex, but this does not allow for direct competition between the two modes of reproduction as the populations are not separable. Instead, one must compare distinct sexual and asexual lineages (e.g. water snails, Lively, 1987; onion thrips, Kobayashi *et al*., 2013), which are relatively rare in nature (possibly because other factors such as mutation accumulation make long-term coexistence unstable; West *et al*., 1999). It is, however, possible to experimentally manipulate some species in the laboratory to obtain distinct sexual and asexual lineages (e.g. nematodes; Morran *et al*., 2011), which may be more amenable to testing the model predictions.

Second, while the model is unable to identify suitable host populations, it does provide some insights as to parasite characteristics that select for sex, which should narrow one side of the search for suitable study species. Specifically, the results suggest which epidemiological characteristics are likely to be the most important for the maintenance of sex (intermediate mortality virulence, higher sterility virulence, moderate to high transmissibility, and moderate to low recovery) and hence which infectious agents may be the best targets for testing whether sex is maintained by parasitism. Interestingly, a water snail-trematode system, which is probably the best-studied empirical example of the maintenance of sex by parasites, involves castration of the host (Lively, 1987). One might speculate from the results of the present study – where sterility virulence increases selection for sex provided there are no additional costs to mating – that castration is a key feature making this particular parasite (as opposed to other parasites of this host which do not castrate) crucial for maintaining host sex. By targeting future searches for study systems towards parasites with the epidemiological characteristics identified in the present study, one may avoid focusing on parasites that are unlikely to affect selection for sex.

In summary, the modelling herein has shown when parasitism maintains sex in the absence of RQD, and how interactions between host demography and parasite epidemiology affect the frequency of sex in the host population. Since most previous studies neither incorporate ecological dynamics nor consider stable coexistence between sex and asex, the present study addresses a major gap in the literature. Mortality virulence appears to be critical in determining selection for or against sex, as it interacts strongly with all parameters and mediates both overall disease prevalence and the cost of being infected. The model also reveals that pairwise interactions between parameters can lead to substantial variation in the frequency of sex, and therefore the interplay between demographic and epidemiological processes is likely to play a major role in the maintenance of sex by parasites.

## Acknowledgements

This work was supported by the Natural Environment Research Council (grant number NE/N014979/1).

## Data availability

Source code is available in the following Github repository: https://github.com/ecoevogroup/Ashby_parasitism_sex_2020.

