## Supplementary material for "When does parasitism maintain sex in the absence of Red Queen Dynamics?": Source code and data: convertToHaplotypeFreqs.m

```
function haplotypeFreqs = convertToHaplotypeFreqs(input,haplotypes,genotypes,n)

% Converts ODE output data to haplotype frequencies

% Normalise input
input = input./sum(input,2);

haplotypeFreqs = zeros(length(input(:,1)),length(haplotypes(:,1)));


for i=1:length(haplotypes(:,1))
    list = find(sum(haplotypes(i,:)==genotypes(:,1:n),2)==n);
    
    haplotypeFreqs(:,i) = sum(input(:,list),2);
end
```
