## Supplementary material for "When does parasitism maintain sex in the absence of Red Queen Dynamics?": Source code and data: fig_dxd_disprev.m

```
function fig_dxd_disprev

% Generates Fig. 3 - demographic x demographic parameters (disease
% prevalence)

% Fixed values
RES1 = 101;
RES2 = 100;
mx = 1;

% Figure dimensions - may differ depending on display
figure(3)
clf
set(gcf,'color','w')
set(gcf,'PaperUnits','centimeters')
xSize = 16; ySize = 5;
xLeft = (21-xSize)/2; yTop = (30-ySize)/2;
set(gcf,'PaperPosition',[xLeft yTop xSize ySize])
set(gcf,'Position',[10 100 xSize*50 ySize*50])

panelcount = 0;
for i1=1:2
    if(i1==1)
        str1 = 'b';
        s1 = 'birth rate, $b$';
    else
        str1 = 'd';
        s1 = 'death rate, $d$';
    end
    for j1=1:2
        if(j1==1)
            str2 = 'd';
            s2 = 'death rate, $d$';
        else
            str2 = 'q';
            s2 = 'competition, $q$';
        end
        filename = strcat('Data/heatmapSweepFinal_',str1,'_',str2,'_',num2str(RES1),'_',num2str(RES2),'.mat');
        if(exist(filename,'file'))
            load(filename)
            DISPREV_MEAN_FULL=[];
            for k1=1:asex_strain_total
                DISPREV_MEAN_FULL = cat(3,DISPREV_MEAN_FULL,repmat(DISPREV_MEAN(:,:,k1),[1,1,host_genotypes_grouped(k1,2)]));
            end
            panelcount = panelcount+1;
            subplot(1,3,panelcount)
            contourf(V2,V1,mean(DISPREV_MEAN_FULL,3),0:0.1:1);
            
            set(gca,'fontsize',10)
            ylabel(s1,'interpreter','latex','fontsize',14)
            xlabel(s2,'interpreter','latex','fontsize',14)
            set(gca,'yscale','log')
            set(gca,'xscale','log')
            
            view(2)
            box on
            set(gca,'xgrid','off')
            set(gca,'ygrid','off')
            
            if(panelcount==2)
                T=title('\bf{demographic $\times$ demographic parameters}','fontsize',16,'interpreter','latex');
                temp = get(T,'position');
                temp(2) = temp(2) + 7;
                set(T,'position',temp)
            end
        end
    end
end
drawnow
C = colorbar('eastoutside');
temp = get(C,'position');
temp(1) = temp(1) + 0.05;
temp(3) = temp(3) + 0.01;
temp(4) = temp(4) - 0.1;
set(C,'position',temp)
set(C,'ytick',0:0.2:mx)
y2=ylabel(C,'disease prevalence','interpreter','latex','fontsize',14);

labs = {'(a)','(b)','(c)'};
textpos = [0.09,0.15
    0.00035,0.15
    0.00035,0.7];
for panelcount=1:3
    subplot(1,3,panelcount)
    set(gca,'clim',[0,mx])
    temp(panelcount,:)=get(gca,'position');
    temp(panelcount,1) = temp(panelcount,1)-0.05;
    temp(panelcount,4) = temp(panelcount,4)-0.1;
    if(panelcount==3)
        temp(panelcount,3) = temp(1,3);
    end
        
    set(gca,'position',temp(panelcount,:))
    x1 = get(gca,'xlim');
    y1 = get(gca,'ylim');
    text(x1(1),y1(end)*1.3,strcat('\bf{',labs{panelcount},'}'),'fontsize',12,'interpreter','latex')
    text(textpos(panelcount,1),textpos(panelcount,2),'sex unviable','interpreter','latex')
end
drawnow
temp = get(y2,'position');
temp(1) = temp(1)-4;
set(y2,'position',temp)

colormap('hot')

save2pdf('fig_dxd_disprev.pdf',gcf,1200)
```
