## Supplementary material for "When does parasitism maintain sex in the absence of Red Queen Dynamics?": Source code and data: fig_dxe_disprev.m

```
function fig_dxe_disprev

% Generates Fig. 7 - demographic x epidemiological parameters (disease
% prevalence)

% Fixed values
RES1 = 101;
RES2 = 100;
mx = 1;

% Figure dimensions - may differ depending on display
figure(7)
clf
set(gcf,'color','w')
set(gcf,'PaperUnits','centimeters')
xSize = 16; ySize = 12;
xLeft = (21-xSize)/2; yTop = (30-ySize)/2;
set(gcf,'PaperPosition',[xLeft yTop xSize ySize])
set(gcf,'Position',[10 100 xSize*50 ySize*50])

panelcount = 0;
for i1=1:3
    if(i1==1)
        str1 = 'b';
        s1 = 'birth rate, $b$';
    elseif(i1==2)
        str1 = 'd';
        s1 = 'death rate, $d$';
    else
        str1 = 'q';
        s1 = 'competition, $q$';
    end
    for j1=1:4
        if(j1==1)
            str2 = 'alpha';
            s2 = {'mortality','virulence, $\alpha$'};
        elseif(j1==2)
            str2 = 'f';
            s2 = {'sterility','virulence, $f$'};
        elseif(j1==3)
            str2 = 'beta';
            s2 = {'transmission','rate, $\beta$'};
        else
            str2 = 'gamma';
            s2 = {'recovery','rate, $\gamma$'};
        end
        filename = strcat('Data/heatmapSweepFinal_',str1,'_',str2,'_',num2str(RES1),'_',num2str(RES2),'.mat');
        if(exist(filename,'file'))
            load(filename)
            DISPREV_MEAN_FULL=[];
            for k1=1:asex_strain_total
                DISPREV_MEAN_FULL = cat(3,DISPREV_MEAN_FULL,repmat(DISPREV_MEAN(:,:,k1),[1,1,host_genotypes_grouped(k1,2)]));
            end
            panelcount = panelcount+1;
            subplot(3,4,panelcount)
            
            contourf(V2,V1,mean(DISPREV_MEAN_FULL,3),0:0.1:mx);
            set(gca,'clim',[0,mx])
            
            set(gca,'fontsize',10)
            if(j1==1)
                ylabel(s1,'interpreter','latex','fontsize',14)
            end
            
            if(i1<3)
                set(gca,'xtick',[])
            else
                if(j1==1)
                    set(gca,'xtick',10.^[-2,0,2])
                elseif(j1==4)
                    set(gca,'xtick',10.^[-2,-1,0,1])
                end
            end
            if(j1>1)
                set(gca,'ytick',[])
            end
            
            if(i1==3)
                Xlab(j1)=xlabel(s2,'interpreter','latex','fontsize',14);
            end
            set(gca,'yscale','log')
            if(j1~=2)
                set(gca,'xscale','log')
            end
            
            view(2)
            box on
            set(gca,'xgrid','off')
            set(gca,'ygrid','off')
        end
    end
end
drawnow
C = colorbar('eastoutside');
temp = get(C,'position');
temp(1) = temp(1) + 0.06;
temp(3) = temp(3) + 0.02;
temp(4) = temp(4) + 0.6;
set(C,'position',temp)
set(C,'ytick',0:0.2:1)
y2=ylabel(C,'disease prevalence','interpreter','latex','fontsize',14);

labs = {'(a.i)','(a.ii)','(a.iii)','(a.iv)','(b.i)','(b.ii)','(b.iii)','(b.iv)','(c.i)','(c.ii)','(c.iii)','(c.iv)'};
textpos = [0.05,0.14
    1/4,0.14
    0.03,0.14
    0.05,0.14
    0.05,0.68
    1/4,0.68
    0.03,0.68
    0.05,0.68
    NaN,NaN
    NaN,NaN
    NaN,NaN
    NaN,NaN];
temp=[];
for panelcount=1:12
    subplot(3,4,panelcount)
    temp(panelcount,:)=get(gca,'position');
    temp(panelcount,1) = temp(panelcount,1)-0.05;
    if(panelcount<5)
        temp(panelcount,2) = temp(panelcount,2)-0.02;
    end
    if(panelcount>4 && panelcount<9)
        temp(panelcount,2) = temp(panelcount,2)-0.01;
    end
    temp(panelcount,3) = temp(panelcount,3)+0.02;
    temp(panelcount,4) = temp(panelcount,4)+0.02;
    set(gca,'position',temp(panelcount,:))
    x1 = get(gca,'xlim');
    y1 = get(gca,'ylim');
    text(x1(1),y1(end)*1.3,strcat('\bf{',labs{panelcount},'}'),'fontsize',12,'interpreter','latex')
    text(textpos(panelcount,1),textpos(panelcount,2),'sex unviable','interpreter','latex')
    
    if(panelcount==2)
        T=title('\bf{demographic $\times$ epidemiological parameters}','fontsize',16,'interpreter','latex');
        temp2 = get(T,'position');
        temp2(1) = temp2(1) + 0.8;
        temp2(2) = temp2(2) + 9;
        set(T,'position',temp2)
    end
end
temp = get(y2,'position');
temp(1) = temp(1)-3.2;
set(y2,'position',temp)

for i=1:4
    temp=get(Xlab(i),'position');
    temp(2) = 0.4*1e-4;
    if(i==4)
        temp(2) = 0.34*1e-4;
    end
    set(Xlab(i),'position',temp)
end

colormap('hot')

save2pdf('fig_dxe_disprev.pdf',gcf,1200)
```
