## Supplementary material for "When does parasitism maintain sex in the absence of Red Queen Dynamics?": Source code and data: fig_dynamics.m

```
function fig_dynamics

% Generates Fig. 1 - example dynamics

% Parameters
t_max = 1e2;
b = 1;
d = 0.1;
q = 1e-3;
alpha = 1;
f = 0.5;
beta = 0.1;
gamma = 0.1; % Note different to default to show transient RQD
eqtol = 0;
c = 0.5;
r = 0.1;

% Figure labels
labs = {'(a.i)','(b.i)','(c.i)'};
labs2 = {'(a.ii)','(b.ii)','(c.ii)'};
labs3 = {'(a.iii)','(b.iii)','(c.iii)'};

% Figure dimensions - may differ depending on display
figure(1)
clf
set(gcf,'color','w')
set(gcf,'PaperUnits','centimeters')
xSize = 16; ySize = 10;
xLeft = (21-xSize)/2; yTop = (30-ySize)/2;
set(gcf,'PaperPosition',[xLeft yTop xSize ySize])
set(gcf,'Position',[10 100 xSize*50 ySize*50])

% Loop over different number of loci
for n=1:3    
    % Number of host and parasite genotypes
    host_geno_total = 2^(2*n);
    par_geno_total = 2^n;
    
    % Set up host and parasite genetics, either dynamically or through a
    % presaved look-up table (LUT).
    % NB. The recombination matrix gives the proportion of each haplotype 
    % (columns) produced by each host genotype (rows) when the recombination
    % rate between each locus is r. 
    if(~exist(strcat('Data/LUT_',num2str(n),'_',num2str(r),'.mat'),'file'))
        [haplotypes,host_genotypes,parasite_genotypes,recombination_matrix,host_genotypes_grouped] = genetics(n,r);
        save(strcat('Data/LUT_',num2str(n),'_',num2str(r),'.mat'),'haplotypes','host_genotypes','parasite_genotypes','recombination_matrix','host_genotypes_grouped')
    else
        load(strcat('Data/LUT_',num2str(n),'_',num2str(r),'.mat'))
    end
    asex_strain_total = length(host_genotypes_grouped(:,1)); % This is actually the number of different types of genotypes based on permutations of the haplotypes
    
    % Infection genetics
    Q = zeros(host_geno_total,par_geno_total);
    for i=1:host_geno_total
        for j=1:par_geno_total
            for k=1:2
                Q(i,j) = Q(i,j) + all(host_genotypes(i,((k-1)*n + 1):(k*n))==parasite_genotypes(j,:))/2;
            end
        end
    end
    
    % Initial conditions
    if(b*c-d>0)
        if((beta*((b*c-d)/(b*c*q))/((d+alpha+gamma)*(2^n)))>1)
            S = (d+alpha+gamma)*(2^n)/beta;
            GAMMA = (d+alpha+gamma);
            if(f>0)
                I = [-(GAMMA*2^n*b*c*f*q + GAMMA*2^n*b*c*q - b*beta*c*f + GAMMA*beta - gamma*beta - sqrt(GAMMA^2*(2^n)^2*b^2*c^2*f^2*q^2 + 2*GAMMA^2*(2^n)^2*b^2*c^2*f*q^2 + GAMMA^2*(2^n)^2*b^2*c^2*q^2 - 4*GAMMA^2*4^n*b^2*c^2*f*q^2 - 2*GAMMA*2^n*b^2*beta*c^2*f^2*q + 2*GAMMA*2^n*b^2*beta*c^2*f*q + 2*GAMMA^2*2^n*b*beta*c*f*q - 2*GAMMA*2^n*gamma*b*beta*c*f*q - 4*GAMMA*2^n*b*beta*c*d*f*q + b^2*beta^2*c^2*f^2 + 2*GAMMA^2*2^n*b*beta*c*q - 2*GAMMA*2^n*gamma*b*beta*c*q - 2*GAMMA*b*beta^2*c*f + 2*gamma*b*beta^2*c*f + GAMMA^2*beta^2 - 2*GAMMA*gamma*beta^2 + gamma^2*beta^2))/(2*b*c*f*q*beta), -(GAMMA*2^n*b*c*f*q + GAMMA*2^n*b*c*q - b*beta*c*f + GAMMA*beta - gamma*beta + sqrt(GAMMA^2*(2^n)^2*b^2*c^2*f^2*q^2 + 2*GAMMA^2*(2^n)^2*b^2*c^2*f*q^2 + GAMMA^2*(2^n)^2*b^2*c^2*q^2 - 4*GAMMA^2*4^n*b^2*c^2*f*q^2 - 2*GAMMA*2^n*b^2*beta*c^2*f^2*q + 2*GAMMA*2^n*b^2*beta*c^2*f*q + 2*GAMMA^2*2^n*b*beta*c*f*q - 2*GAMMA*2^n*gamma*b*beta*c*f*q - 4*GAMMA*2^n*b*beta*c*d*f*q + b^2*beta^2*c^2*f^2 + 2*GAMMA^2*2^n*b*beta*c*q - 2*GAMMA*2^n*gamma*b*beta*c*q - 2*GAMMA*b*beta^2*c*f + 2*gamma*b*beta^2*c*f + GAMMA^2*beta^2 - 2*GAMMA*gamma*beta^2 + gamma^2*beta^2))/(2*b*c*f*q*beta)];
                I = I(I>0);
            else
                I=-GAMMA*2^n*(GAMMA*2^n*b*c*q - b*beta*c + beta*d)/(beta*(GAMMA*2^n*b*c*q + GAMMA*beta - gamma*beta));
            end
        else
            S = (b*c-d)/(b*c*q);
            I = 0;
        end
    else
        S=0;
        I=0;
    end
    S_sex = ones(host_geno_total,1)*S/host_geno_total;
    I_sex = ones(host_geno_total,par_geno_total)*I/(host_geno_total*par_geno_total);
    S_asex = zeros(host_geno_total,1);
    I_asex = zeros(host_geno_total,par_geno_total);
    init_pop = [S_sex;S_asex;sum(I_sex,2);sum(I_asex,2);sum(I_sex,1)';sum(I_asex,1)']';
    
    if(S>0)
        if(I==0) % Introduce disease at a low level if it is absent
            I_sex = ones(host_geno_total,par_geno_total)*S/(host_geno_total*par_geno_total*100);
            init_pop = [S_sex;S_asex;sum(I_sex,2);sum(I_asex,2);sum(I_sex,1)';sum(I_asex,1)']';
        end
        
        % Repeated haplotype
        asex_strain = 1;
        init_pop_new = init_pop;
        init_pop_new(host_geno_total + host_genotypes_grouped(asex_strain,1)) = S/1000;
        [t1,S_sex,S_asex,IH_sex,IH_asex,IP_sex,IP_asex] = sexVsAsexSolverFinal(t_max,b,c,d,f,q,alpha,beta,gamma,host_geno_total,par_geno_total,eqtol,init_pop_new,recombination_matrix,Q);
        X1 = [S_sex,S_asex,IH_sex,IH_asex,IP_sex,IP_asex];
        SEX1 = sum(X1(:,[1:host_geno_total,(2*host_geno_total+1):(3*host_geno_total)]),2)./sum(X1(:,1:(4*host_geno_total)),2);
        hF_sex1 = convertToHaplotypeFreqs(S_sex+IH_sex,haplotypes,host_genotypes,n);
        
        % Non-repeated haplotype
        asex_strain = ceil(asex_strain_total/2);
        init_pop_new = init_pop;
        init_pop_new(host_geno_total + host_genotypes_grouped(asex_strain,1)) = S/1000;
        [t2,S_sex,S_asex,IH_sex,IH_asex,IP_sex,IP_asex] = sexVsAsexSolverFinal(t_max,b,c,d,f,q,alpha,beta,gamma,host_geno_total,par_geno_total,eqtol,init_pop_new,recombination_matrix,Q);
        X2 = [S_sex,S_asex,IH_sex,IH_asex,IP_sex,IP_asex];
        SEX2 = sum(X2(:,[1:host_geno_total,(2*host_geno_total+1):(3*host_geno_total)]),2)./sum(X2(:,1:(4*host_geno_total)),2);
        hF_sex2 = convertToHaplotypeFreqs(S_sex+IH_sex,haplotypes,host_genotypes,n);
        
        subplot(2,3,n)
        hold on
        plot(t1,SEX1,'k','linewidth',2)
        plot(t2,SEX2,'color',0.67*[1,1,1],'linewidth',2)
        
        set(gca,'fontsize',10)
        text(0,1.05,strcat('\bf{',labs{n},'}'),'fontsize',12,'interpreter','latex')
        title(strcat('$n=',num2str(n),'$'),'fontsize',12,'interpreter','latex')
        ylim([0,1])
        if(n==1)
            text(0.54*t_max,SEX1(end)+0.05,'homozygous','fontsize',10,'interpreter','latex')
            text(0.54*t_max,SEX2(end)+0.05,'heterozygous','fontsize',10,'interpreter','latex')
            ylabel('Frequency of sex','interpreter','latex','fontsize',14)
        end
        box on
        
        subplot(4,3,6+n)
        hold on
        plot(t1,hF_sex1,'linewidth',1.5)
        plot(t1,1-SEX1,'k:','linewidth',1.5)
        set(gca,'fontsize',10)
        
        ylim([0,max(ceil(10*max(max(hF_sex1(:)),max(hF_sex2(:)))))/10])
        temp=get(gca,'ylim');
        text(0,1.1*temp(2),strcat('\bf{',labs2{n},'}'),'fontsize',12,'interpreter','latex')
        title('homozygous','fontsize',10,'interpreter','latex')
        set(gca,'ytick',0:(max(temp(2)/2,0.1)):temp(2))
        box on        
        
        subplot(4,3,9+n)
        hold on
        plot(t2,hF_sex2,'linewidth',1.5)
        plot(t2,1-SEX2,'k:','linewidth',1.5)
        set(gca,'fontsize',10)
        
        ylim([0,max(ceil(10*max([max(hF_sex1(:)),max(hF_sex2(:)),max(1-SEX1),max(1-SEX2)])))/10])
        temp=get(gca,'ylim');
        text(0,1.1*temp(2),strcat('\bf{',labs3{n},'}'),'fontsize',12,'interpreter','latex')
        set(gca,'ytick',0:(max(temp(2)/2,0.1)):temp(2))
        title('heterozygous','fontsize',10,'interpreter','latex')
        if(n==1)
            y1=ylabel('Haploype frequency','interpreter','latex','fontsize',14);
            temp=get(y1,'position');
            temp(2)=temp(2)+0.8;
            set(y1,'position',temp)
        elseif(n==2)
            xlabel('Time','interpreter','latex','fontsize',14)
        end
        box on
        temp=get(gca,'position');
        temp(2)=temp(2)-0.02;
        set(gca,'position',temp)
        
    end
end

save2pdf('fig_dynamics.pdf')
```
