## Supplementary material for "When does parasitism maintain sex in the absence of Red Queen Dynamics?": Source code and data: fig_exe_disprev.m

```
function fig_exe_disprev

% Generates Fig. 5 - epidemiological x epidemiological parameters (disease
% prevalence)

% Fixed values
RES1 = 101;
RES2 = 100;
mx = 1;

% Figure dimensions - may differ depending on display
figure(5)
clf
set(gcf,'color','w')
set(gcf,'PaperUnits','centimeters')
xSize = 16; ySize = 9;
xLeft = (21-xSize)/2; yTop = (30-ySize)/2;
set(gcf,'PaperPosition',[xLeft yTop xSize ySize])
set(gcf,'Position',[10 100 xSize*50 ySize*50])

for panelcount=1:6
    if(panelcount==1)
        str1 = 'alpha';
        s1 = {'mortality','virulence, $\alpha$'};
        str2 = 'f';
        s2 = {'sterility','virulence, $f$'};
    elseif(panelcount==2)
        str1 = 'alpha';
        s1 = {'mortality','virulence, $\alpha$'};
        str2 = 'beta';
        s2 = {'transmission','rate, $\beta$'};
    elseif(panelcount==3)
        str1 = 'alpha';
        s1 = {'mortality','virulence, $\alpha$'};
        str2 = 'gamma';
        s2 = {'recovery','rate, $\gamma$'};
    elseif(panelcount==4)
        str1 = 'f';
        s1 = {'sterility','virulence, $f$'};
        str2 = 'beta';
        s2 = {'transmission','rate, $\beta$'};
    elseif(panelcount==5)
        str1 = 'f';
        s1 = {'sterility','virulence, $f$'};
        str2 = 'gamma';
        s2 = {'recovery','rate, $\gamma$'};
    else
        str1 = 'beta';
        s1 = {'transmission','rate, $\beta$'};
        str2 = 'gamma';
        s2 = {'recovery','rate, $\gamma$'};
    end
    filename = strcat('Data/heatmapSweepFinal_',str1,'_',str2,'_',num2str(RES1),'_',num2str(RES2),'.mat');
    if(exist(filename,'file'))
        load(filename)
        DISPREV_MEAN_FULL=[];
        for k1=1:asex_strain_total
            DISPREV_MEAN_FULL = cat(3,DISPREV_MEAN_FULL,repmat(DISPREV_MEAN(:,:,k1),[1,1,host_genotypes_grouped(k1,2)]));
        end
        subplot(2,3,panelcount)
        
        contourf(V2,V1,mean(DISPREV_MEAN_FULL,3),0:0.1:1,'linecolor','k');
        set(gca,'clim',[0,mx])
        
        set(gca,'fontsize',10)
        ylabel(s1,'interpreter','latex','fontsize',14)
        Xlab(panelcount)=xlabel(s2,'interpreter','latex','fontsize',14);
        
        if(panelcount<4)
            set(gca,'ytick',10.^(-2:2:2))
        end
        if(panelcount==3 || panelcount==5 || panelcount==6)
            set(gca,'xtick',10.^[-2,-1,0,1])
        end
        
        if(panelcount>1)
            set(gca,'xscale','log')
        end
        
        if(panelcount~=4 && panelcount~=5)
            set(gca,'yscale','log')
        end
        
        view(2)
        box on
        set(gca,'xgrid','off')
        set(gca,'ygrid','off')        
    end
end

drawnow
C = colorbar('eastoutside');
temp = get(C,'position');
temp(1) = temp(1) + 0.06;
temp(2) = temp(2) - 0.03;
temp(3) = temp(3) + 0.02;
temp(4) = temp(4) + 0.49;
set(C,'position',temp)
set(C,'ytick',0:0.2:1)
y2=ylabel(C,'disease prevalence','interpreter','latex','fontsize',14);

labs = {'(a)','(b)','(c)','(d)','(e)','(f)'};
temp=[];
for panelcount=1:6
    subplot(2,3,panelcount)
    temp(panelcount,:)=get(gca,'position');
    temp(panelcount,1) = temp(panelcount,1)-0.05;
    temp(panelcount,2) = temp(panelcount,2)-0.03;
    temp(panelcount,3) = temp(panelcount,3)+0.0;
    temp(panelcount,4) = temp(panelcount,4)+0.02;
    
    set(gca,'position',temp(panelcount,:))
    x1 = get(gca,'xlim');
    y1 = get(gca,'ylim');
    
    if(panelcount<4)
        text(x1(1),y1(end)*1.9,strcat('\bf{',labs{panelcount},'}'),'fontsize',12,'interpreter','latex')
    elseif(panelcount==4 || panelcount==5)
        text(x1(1),y1(end)*1.0704,strcat('\bf{',labs{panelcount},'}'),'fontsize',12,'interpreter','latex')
    else
        text(x1(1),y1(end)*1.5,strcat('\bf{',labs{panelcount},'}'),'fontsize',12,'interpreter','latex')
    end
    
    if(panelcount==2)
        T=title('\bf{epidemiological $\times$ epidemiological parameters}','fontsize',16,'interpreter','latex');
        temp2 = get(T,'position');
        temp2(2) = temp2(2) + 140;
        set(T,'position',temp2)
    end
end
temp = get(y2,'position');
temp(1) = temp(1)-3.2;
set(y2,'position',temp)

drawnow
for i=1:6
    temp=get(Xlab(i),'position');
    if(i==3)
        temp(2) = 0.1*1e-2;
    end
    if(i==5)
        temp(2) = -0.24;
    end
    if(i==6)
        temp(2) = 0.33*1e-2;
    end
    set(Xlab(i),'position',temp)
end

colormap('hot')

save2pdf('fig_exe_disprev.pdf',gcf,1200)
```
