## Supplementary material for "When does parasitism maintain sex in the absence of Red Queen Dynamics?": Source code and data: figGen.m

```
function figGen

% Generates manuscript figures

fig_dynamics % Fig. 1
fig_dxd_freq_sex % Fig. 2
fig_dxd_disprev % Fig. 3
fig_exe_freq_sex % Fig. 4
fig_exe_disprev % Fig. 5
fig_dxe_freq_sex % Fig. 6
fig_dxe_disprev % Fig. 7
```
