## Supplementary material for "When does parasitism maintain sex in the absence of Red Queen Dynamics?": Source code and data: genetics.m

```
function [haplotypes,host_genotypes,parasite_genotypes,recombination_matrix,host_genotypes_grouped] = genetics(n,r)

% Returns genetic data for use in ODE solver

% Define genetics
haplotypes = de2bi(0:(2^n-1));
parasite_genotypes = haplotypes;
temp0 = bi2de(haplotypes);

host_genotypes = [];
for i=1:(2^n)
    host_genotypes = [host_genotypes;repmat(haplotypes(i,:),[2^n,1]),haplotypes];
end
host_genotypes = sortrows(host_genotypes,(2*n):-1:1);
recombination_matrix = zeros(2^(2*n),2^n);

C0 = {};
count1 = 2;
for k=1:(n-1)
    C = combnk(1:(n-1),k); % This gives where the recombination events occur
    C = [C,n*ones(length(C(:,1)),1)];
    count1 = count1 + 2*length(C(:,1));
    C0{k} = C;
end

for i=1:(2^(2*n))
    % This routine gives the pool of haplotypes from host genotype i
    % (no mutation) due to recombination and their associated
    % probabilities
    hap1 = host_genotypes(i,1:n);
    hap2 = host_genotypes(i,(n+1):(2*n));
    possible_haplotypes = zeros(count1,n);
    prob_list_temp = zeros(count1,1);
    possible_haplotypes(1:2,:) = [hap1;hap2];
    prob_list_temp(1:2) = [(1-r)^(n-1);(1-r)^(n-1)];
    count2 = 3;
    for k=1:(n-1)
        C = C0{k};
        
        % Work through rows - different haplotypes
        for j=1:length(C(:,1))
            hap_temp1 = hap1(1);
            hap_temp2 = hap2(1);
            
            % Work through columns - create pairs of haplotypes
            orig = 1;
            curr = 1;
            for l=2:n
                if(l>C(j,curr))
                    orig = mod(orig+1,2);
                    curr=curr+1;
                end
                if(orig>0)
                    hap_temp1(l) = hap1(l);
                    hap_temp2(l) = hap2(l);
                else
                    hap_temp1(l) = hap2(l);
                    hap_temp2(l) = hap1(l);
                end
            end
            possible_haplotypes(count2:(count2+1),:) = [hap_temp1;hap_temp2];
            prob_list_temp(count2:(count2+1)) = [r^(length(C(j,:))-1)*((1-r)^(n-length(C(j,:))));r^(length(C(j,:))-1)*((1-r)^(n-length(C(j,:))))];
            count2 = count2+2;
        end
    end
    prob_list_temp = prob_list_temp/sum(prob_list_temp); % Normalise
    haplotype_list = unique(possible_haplotypes,'rows'); % Find unique set of haplotypes
    temp1 = bi2de(possible_haplotypes);
    temp2 = bi2de(haplotype_list);
    for k=1:length(haplotype_list(:,1))
        temp_list1 = temp1==temp2(k);
        temp_list2 = temp0==temp2(k);
        prob = sum(prob_list_temp(temp_list1));
        recombination_matrix(i,temp_list2) = prob;
    end
    recombination_matrix(i,:) = recombination_matrix(i,:)/sum(recombination_matrix(i,:)); % Normalise
end

% Group genotypes for more efficient parameter sweeps
list1 = sum(host_genotypes,2);
list2 = unique(list1);

for i=1:length(list2)
    list3(i,1) = find(list1==list2(i),1);
    count(i,1) = length(find(list1==list2(i)));
end

host_genotypes_grouped = sortrows([list3,count]);
```
