## Supplementary material for "When does parasitism maintain sex in the absence of Red Queen Dynamics?": Source code and data: heatmapSweepFinal.m

```
function heatmapSweepFinal(params,RES1,RES2)

% Generates data for Fig. 2-7
% Note: params = [b,d,q,alpha,f,beta,gamma];

% Fixed parameters
t_max = 5000;
start = t_max*0.8;
eqtol = 1e-2;
c = 0.5;
n = 3;
r = 0.1;

% Parse inputs
[V1,V2,str1,str2] = parseInputs(params,RES1,RES2);
p1 = find(isnan(params),1);
p2 = find(isnan(params),1,'last');
b = params(1);
d = params(2);
q = params(3);
alpha = params(4);
f = params(5);
beta = params(6);
gamma = params(7);


% Set output filename and check if file already exists
filename  = strcat('Data/heatmapSweepFinal_',str1,'_',str2,'_',num2str(RES1),'_',num2str(RES2),'.mat');
if(exist(filename,'file'))
    disp(strcat('skipping:',filename));
else
    
    % Number of host and parasite genotypes
    host_geno_total = 2^(2*n);
    par_geno_total = 2^n;
    
    % Set up host and parasite genetics, either dynamically or through a
    % presaved look-up table (LUT).
    % NB. The recombination matrix gives the proportion of each haplotype 
    % (columns) produced by each host genotype (rows) when the recombination
    % rate between each locus is r. 
    if(~exist(strcat('Data/LUT_',num2str(n),'_',num2str(r),'.mat'),'file'))
        [haplotypes,host_genotypes,parasite_genotypes,recombination_matrix,host_genotypes_grouped] = genetics(n,r);
        save(strcat('Data/LUT_',num2str(n),'_',num2str(r),'.mat'),'haplotypes','host_genotypes','parasite_genotypes','recombination_matrix','host_genotypes_grouped')
    else
        load(strcat('Data/LUT_',num2str(n),'_',num2str(r),'.mat'))
    end
    asex_strain_total = length(host_genotypes_grouped(:,1)); % This is actually the number of different types of genotypes based on permutations of the haplotypes
    
    % Infection genetics
    Q = zeros(host_geno_total,par_geno_total);
    for i=1:host_geno_total
        for j=1:par_geno_total
            for k=1:2
                Q(i,j) = Q(i,j) + all(host_genotypes(i,((k-1)*n + 1):(k*n))==parasite_genotypes(j,:))/2;
            end
        end
    end
    
    % Outputs
    S_INITIAL = NaN*zeros(length(V1),length(V2));
    I_INITIAL = NaN*zeros(length(V1),length(V2));
    N_MEAN = NaN*zeros(length(V1),length(V2),asex_strain_total);
    N_VAR = NaN*zeros(length(V1),length(V2),asex_strain_total);
    N_MIN = NaN*zeros(length(V1),length(V2),asex_strain_total);
    SEX_MEAN = NaN*zeros(length(V1),length(V2),asex_strain_total);
    SEX_VAR = NaN*zeros(length(V1),length(V2),asex_strain_total);
    SEX_MIN = NaN*zeros(length(V1),length(V2),asex_strain_total);
    DISPREV_MEAN = NaN*zeros(length(V1),length(V2),asex_strain_total);
    DISPREV_VAR = NaN*zeros(length(V1),length(V2),asex_strain_total);
    DISPREV_MIN = NaN*zeros(length(V1),length(V2),asex_strain_total);
    DISPREV_SEX_MEAN = NaN*zeros(length(V1),length(V2),asex_strain_total);
    DISPREV_SEX_VAR = NaN*zeros(length(V1),length(V2),asex_strain_total);
    DISPREV_SEX_MIN = NaN*zeros(length(V1),length(V2),asex_strain_total);
    DISPREV_ASEX_MEAN = NaN*zeros(length(V1),length(V2),asex_strain_total);
    DISPREV_ASEX_VAR = NaN*zeros(length(V1),length(V2),asex_strain_total);
    DISPREV_ASEX_MIN = NaN*zeros(length(V1),length(V2),asex_strain_total);
    SEX_GENOTYPES_VAR = NaN*zeros(length(V1),length(V2),asex_strain_total);
    ASEX_GENOTYPES_VAR = NaN*zeros(length(V1),length(V2),asex_strain_total);
    
    % Loop over parameter 2 vector
    TOTAL = length(V2);
    COUNT = 0;
    for j=1:length(V2)
        if(p2==1)
            b = V2(j);
        elseif(p2==2)
            d = V2(j);
        elseif(p2==3)
            q = V2(j);
        elseif(p2==4)
            alpha = V2(j);
        elseif(p2==5)
            f = V2(j);
        elseif(p2==6)
            beta = V2(j);
        else
            gamma = V2(j);
        end
        tic;
        % Loop over parameter 1 vector
        for i=1:length(V1)
            if(p1==1)
                b = V1(i);
            elseif(p1==2)
                d = V1(i);
            elseif(p1==3)
                q = V1(i);
            elseif(p1==4)
                alpha = V1(i);
            elseif(p1==5)
                f = V1(i);
            elseif(p1==6)
                beta = V1(i);
            else
                gamma = V1(i);
            end
            
            % Initial conditions
            if(b*c-d>0)
                if((beta*((b*c-d)/(b*c*q))/((d+alpha+gamma)*(2^n)))>1)
                    S = (d+alpha+gamma)*(2^n)/beta;
                    GAMMA = (d+alpha+gamma);
                    if(f>0)
                        I = [-(GAMMA*2^n*b*c*f*q + GAMMA*2^n*b*c*q - b*beta*c*f + GAMMA*beta - gamma*beta - sqrt(GAMMA^2*(2^n)^2*b^2*c^2*f^2*q^2 + 2*GAMMA^2*(2^n)^2*b^2*c^2*f*q^2 + GAMMA^2*(2^n)^2*b^2*c^2*q^2 - 4*GAMMA^2*4^n*b^2*c^2*f*q^2 - 2*GAMMA*2^n*b^2*beta*c^2*f^2*q + 2*GAMMA*2^n*b^2*beta*c^2*f*q + 2*GAMMA^2*2^n*b*beta*c*f*q - 2*GAMMA*2^n*gamma*b*beta*c*f*q - 4*GAMMA*2^n*b*beta*c*d*f*q + b^2*beta^2*c^2*f^2 + 2*GAMMA^2*2^n*b*beta*c*q - 2*GAMMA*2^n*gamma*b*beta*c*q - 2*GAMMA*b*beta^2*c*f + 2*gamma*b*beta^2*c*f + GAMMA^2*beta^2 - 2*GAMMA*gamma*beta^2 + gamma^2*beta^2))/(2*b*c*f*q*beta), -(GAMMA*2^n*b*c*f*q + GAMMA*2^n*b*c*q - b*beta*c*f + GAMMA*beta - gamma*beta + sqrt(GAMMA^2*(2^n)^2*b^2*c^2*f^2*q^2 + 2*GAMMA^2*(2^n)^2*b^2*c^2*f*q^2 + GAMMA^2*(2^n)^2*b^2*c^2*q^2 - 4*GAMMA^2*4^n*b^2*c^2*f*q^2 - 2*GAMMA*2^n*b^2*beta*c^2*f^2*q + 2*GAMMA*2^n*b^2*beta*c^2*f*q + 2*GAMMA^2*2^n*b*beta*c*f*q - 2*GAMMA*2^n*gamma*b*beta*c*f*q - 4*GAMMA*2^n*b*beta*c*d*f*q + b^2*beta^2*c^2*f^2 + 2*GAMMA^2*2^n*b*beta*c*q - 2*GAMMA*2^n*gamma*b*beta*c*q - 2*GAMMA*b*beta^2*c*f + 2*gamma*b*beta^2*c*f + GAMMA^2*beta^2 - 2*GAMMA*gamma*beta^2 + gamma^2*beta^2))/(2*b*c*f*q*beta)];
                        I = I(I>0);
                    else
                        I=-GAMMA*2^n*(GAMMA*2^n*b*c*q - b*beta*c + beta*d)/(beta*(GAMMA*2^n*b*c*q + GAMMA*beta - gamma*beta));
                    end
                else
                    S = (b*c-d)/(b*c*q);
                    I = 0;
                end
            else
                S=0;
                I=0;
            end
            S_sex = ones(host_geno_total,1)*S/host_geno_total;
            I_sex = ones(host_geno_total,par_geno_total)*I/(host_geno_total*par_geno_total);
            S_asex = zeros(host_geno_total,1);
            I_asex = zeros(host_geno_total,par_geno_total);
            init_pop = [S_sex;S_asex;sum(I_sex,2);sum(I_asex,2);sum(I_sex,1)';sum(I_asex,1)']';
            
            S_INITIAL(i,j) = S;
            I_INITIAL(i,j) = I;
            if(S>0)
                if(I==0) % Introduce disease at a low level if it is absent
                    I_sex = ones(host_geno_total,par_geno_total)*S/(host_geno_total*par_geno_total*100);
                    init_pop = [S_sex;S_asex;sum(I_sex,2);sum(I_asex,2);sum(I_sex,1)';sum(I_asex,1)']';
                end
                % Introduce a single asexual lineage
                for asex_strain = 1:asex_strain_total
                    init_pop_new = init_pop;
                    init_pop_new(host_geno_total + host_genotypes_grouped(asex_strain,1)) = S/1000;
                    
                    % Call ODE solver
                    [t,S_sex,S_asex,IH_sex,IH_asex,IP_sex,IP_asex] = sexVsAsexSolverFinal(t_max,b,c,d,f,q,alpha,beta,gamma,host_geno_total,par_geno_total,eqtol,init_pop_new,recombination_matrix,Q);
                    X = [S_sex,S_asex,IH_sex,IH_asex,IP_sex,IP_asex];
                    
                    T = find(t>start);
                    if(~isempty(T))
                        
                        SEX = sum(X(:,[1:host_geno_total,(2*host_geno_total+1):(3*host_geno_total)]),2)./sum(X(:,1:(4*host_geno_total)),2);
                        DISPREV = sum(X(:,(2*host_geno_total+1):(4*host_geno_total)),2)./sum(X(:,1:(4*host_geno_total)),2);
                        DISPREV_SEX = sum(X(:,(2*host_geno_total+1):(3*host_geno_total)),2)./sum(X(:,[1:host_geno_total,(2*host_geno_total+1):(3*host_geno_total)]),2);
                        DISPREV_ASEX = sum(X(:,(3*host_geno_total+1):(4*host_geno_total)),2)./sum(X(:,[(1+host_geno_total):(2*host_geno_total),(3*host_geno_total+1):(4*host_geno_total)]),2);
                        SEX_GENOTYPES = (X(:,1:host_geno_total) + X(:,(2*host_geno_total+1):(3*host_geno_total)))./repmat(sum((X(:,1:host_geno_total) + X(:,(2*host_geno_total+1):(3*host_geno_total))),2),[1,host_geno_total]);
                        ASEX_GENOTYPES = (X(:,(host_geno_total+1):(2*host_geno_total)) + X(:,(3*host_geno_total+1):(4*host_geno_total)))./repmat(sum((X(:,(host_geno_total+1):(2*host_geno_total)) + X(:,(3*host_geno_total+1):(4*host_geno_total))),2),[1,host_geno_total]);
                        
                        % Population sizes
                        N_MEAN(i,j,asex_strain) = mean(sum(X(T,:),2));
                        N_VAR(i,j,asex_strain) = var(sum(X(T,:),2));
                        N_MIN(i,j,asex_strain) = min(sum(X(T,:),2));
                        
                        % Frequency of sex (mean & var during
                        % measurement window, min overall)
                        SEX_MEAN(i,j,asex_strain) = mean(SEX(T)); % Mean frequency of sex during measurement window
                        SEX_VAR(i,j,asex_strain) = var(SEX(T)); % Variance in frequency of sex during measurement window
                        SEX_MIN(i,j,asex_strain) = min(SEX); % Minimum frequency of sex at any point
                        
                        % Disease prevalence during measurement
                        % window (mean, var, min)
                        DISPREV_MEAN(i,j,asex_strain) = mean(DISPREV(T));
                        DISPREV_VAR(i,j,asex_strain) = var(DISPREV(T));
                        DISPREV_MIN(i,j,asex_strain) = min(DISPREV(T));
                        
                        % Disease prevalence in sexuals during measurement
                        % window (mean, var, min)
                        DISPREV_SEX_MEAN(i,j,asex_strain) = mean(DISPREV_SEX(T));
                        DISPREV_SEX_VAR(i,j,asex_strain) = var(DISPREV_SEX(T));
                        DISPREV_SEX_MIN(i,j,asex_strain) = min(DISPREV_SEX(T));
                        
                        % Disease prevalence in asexuals during measurement
                        % window (mean, var, min)
                        DISPREV_ASEX_MEAN(i,j,asex_strain) = mean(DISPREV_ASEX(T));
                        DISPREV_ASEX_VAR(i,j,asex_strain) = var(DISPREV_ASEX(T));
                        DISPREV_ASEX_MIN(i,j,asex_strain) = min(DISPREV_ASEX(T));
                        
                        % Variance in sexual genotype frequencies
                        % after asexual introduced
                        SEX_GENOTYPES_VAR(i,j,asex_strain) = mean(var(SEX_GENOTYPES(T,:),0,1));
                        
                        % Variance in asexual genotype frequencies
                        % after asexual introduced
                        ASEX_GENOTYPES_VAR(i,j,asex_strain) = mean(var(ASEX_GENOTYPES(T,:),0,1));
                    end
                end
            end
        end
        toc;
        COUNT = COUNT + 1;
        PROGRESS = COUNT/TOTAL
    end
    clear INITIAL i j S_sex S_asex I_sex I_asex t x SEX t1 asex_strain COUNT TOTAL PROGRESS ans t1 S_sex1 S_asex1 IH_sex1 IH_asex1 IP_sex1 IP_asex1 X01 T0
    
    save(filename);
    
end


function [V1,V2,str1,str2] = parseInputs(params,RES1,RES2)

flag = 0;
if(isnan(params(1))) % b
    V1 = logspace(-1,1,RES1);
    flag = 1;
    str1 = 'b';
end

if(isnan(params(2))) % d
    if(flag==0)
        V1 = logspace(-2,0,RES1);
        str1 = 'd';
    else
        V2 = logspace(-2,0,RES2);
        str2 = 'd';
    end
    flag = 1;
end

if(isnan(params(3))) % q
    if(flag==0)
        V1 = logspace(-4,-2,RES1);
        str1 = 'q';
    else
        V2 = logspace(-4,-2,RES2);
        str2 = 'q';
    end
    flag = 1;
end

if(isnan(params(4))) % alpha
    if(flag==0)
        V1 = logspace(-2,2,RES1);
        str1 = 'alpha';
    else
        V2 = logspace(-2,2,RES2);
        str2 = 'alpha';
    end
    flag = 1;
end

if(isnan(params(5))) % f
    if(flag==0)
        V1 = linspace(0,1,RES1);
        str1 = 'f';
    else
        V2 = linspace(0,1,RES2);
        str2 = 'f';
    end
    flag = 1;
end

if(isnan(params(6))) % beta
    if(flag==0)
        V1 = logspace(-2,0,RES1);
        str1 = 'beta';
    else
        V2 = logspace(-2,0,RES2);
        str2 = 'beta';
    end
    flag = 1;
end

if(isnan(params(7))) % gamma
    if(flag==0)
        V1 = logspace(-2,1,RES1);
        str1 = 'gamma';
    else
        V2 = logspace(-2,1,RES2);
        str2 = 'gamma';
    end
end
```
