## Supplementary material for "When does parasitism maintain sex in the absence of Red Queen Dynamics?": Source code and data: heatmapSweepFinalEntry.m

```
function heatmapSweepFinalEntry(RES1,RES2)

% Entry file to generate data for Fig. 2-7

% Default parameters
b = 1;
d = 0.1;
q = 1e-3;
alpha = 1;
f = 0.5;
beta = 0.1;
gamma = 1;
params = [b,d,q,alpha,f,beta,gamma];

% b as lead param
for i=2:7
    params0 = params;
    params0(1) = NaN;
    params0(i) = NaN;
    heatmapSweepFinal(params0,RES1,RES2);
end

% d as lead param
for i=3:7
    params0 = params;
    params0(2) = NaN;
    params0(i) = NaN;
    heatmapSweepFinal(params0,RES1,RES2);
end

% q as lead param
for i=4:7
    params0 = params;
    params0(3) = NaN;
    params0(i) = NaN;
    heatmapSweepFinal(params0,RES1,RES2);
end

% alpha as lead param
for i=5:7
    params0 = params;
    params0(4) = NaN;
    params0(i) = NaN;
    heatmapSweepFinal(params0,RES1,RES2);
end

% f as lead param
for i=6:7
    params0 = params;
    params0(5) = NaN;
    params0(i) = NaN;
    heatmapSweepFinal(params0,RES1,RES2);
end

% beta as lead param
params0 = params; params0(6) = NaN; params0(7) = NaN;
heatmapSweepFinal(params0,RES1,RES2);
```
