## Supplementary material for "When does parasitism maintain sex in the absence of Red Queen Dynamics?": Source code and data: readme.txt

Data and figure generation files for manuscript: When does parasitism maintain sex in the absence of Red Queen Dynamics?
Author: Dr Ben Ashby, University of Bath
Contact:
Notes: All files generated and tested on Malatb 2020a. ODE solver is written as a MEX file in C#. 
Instructions: 1) compile ODE solver in Matlab using mex sexVsAsexSolverFinal.c. 
2) To recreate figures using presaved data, extract 'Data.zip' then call figGen
3) Figure data can be regenerated by calling heatmapSweepFinalEntry followed by figGen

File list:
readme.txt - Readme file
convertToHaplotypeFreqs.m - Converts ODE output data to haplotype frequencies
genetics.m - Returns genetic data for use in ODE solver
heatmapSweepFinal - Generates data for Fig. 2-7
heatmapSweepFinalEntry - Entry file to generate data for Fig. 2-7
colmap.mat - data file for colormap
figGen.m - Generates manuscript figures
fig_dynamics.m - Generates Fig. 1: example dynamics
fig_dxd_freq_sex.m - Generates Fig. 2: demographic x demographic parameters (frequency of sex)
fig_dxd_disprev.m - Generates Fig. 3: demographic x demographic parameters (disease prevalence)
fig_exe_freq_sex.m - Generates Fig. 4: epidemiological x epidemiological parameters (frequency of sex)
fig_exe_disprev.m - Generates Fig. 5: epidemiological x epidemiological parameters (disease prevalence)
fig_dxe_freq_sex.m - Generates Fig. 6: demographic x epidemiological parameters (frequency of sex)
fig_dxe_disprev.m - Generates Fig. 7: demographic x epidemiological parameters (disease prevalence)
Data.zip - Presaved data for figure generation
