## Supplementary material for "When does parasitism maintain sex in the absence of Red Queen Dynamics?": Source code and data: sexVsAsexSolverFinal.c

```
/***********************************************************************************************************
 * [t,S_sex,S_asex,IH_sex,IH_asex,IP_sex,IP_asex] = sexVsAsexSolverFinal(t_max,b,c,d,f,q,alpha,beta,gamma,host_geno_total,par_geno_total,eqtol,init_pop,recombination_matrix,Q);
 ***********************************************************************************************************/

#include <mex.h>
#include <math.h>

/***********************************
 * Constant parameter values
 ***********************************/
#define MAXSTEPS 1e6 /* Maximum number of steps for ODE solver */
#define INTERVAL 1e2 /* Check if the system is close to equilibrium */
#define EPS 1e-6 /* ODE solver tolerance */
#define TINY 1e-30 /* Constant value for solver */
#define TINY2 1e-30 /* Constant value for solver to avoid tolerance issues */
/* RK solver parameters */
#define b21 0.2
#define b31 3.0/40.0
#define b32 9.0/40.0
#define b41 0.3
#define b42 -0.9
#define b43 1.2
#define b51 -11.0/54.0
#define b52 2.5
#define b53 -70.0/27.0
#define b54 35.0/27.0
#define b61 1631.0/55296
#define b62 175.0/512.0
#define b63 575.0/13824.0
#define b64 44275.0/110592
#define b65 253.0/4096.0
#define c1 37.0/378.0
#define c3 250.0/621.0
#define c4 125.0/594.0
#define c6 512.0/1771.0
#define dc5 -277.00/14336

/*************************************
 * Define structure for model parameters
 *************************************/
struct PARAM{
    double t_max;
    double b;
    double c;
    double d;
    double f;
    double q; 
    double alpha;
    double beta;
    double gamma;
    double eqtol;
    int host_geno_total;
    int par_geno_total;
};

/*************************************
 * Function prototypes
 *************************************/
int my_rungkut (double *T, double *S_sex_out, double *S_asex_out, double *IH_sex_out, double *IH_asex_out, double *IP_sex_out, double *IP_asex_out, double *EQFLAG, double *init_pop, double *recombination_matrix, double *Q, struct PARAM *p);
void rkqs(double *S_sex, double *S_asex, double *IH_sex, double *IH_asex, double *IP_sex, double *IP_asex, double *DSSDT, double *DSADT, double *DIHSDT, double *DIHADT, double *DIPSDT, double *DIPADT, double *h, double *hnext, double *SS_SCALE, double *SA_SCALE, double *IHS_SCALE, double *IHA_SCALE, double *IPS_SCALE, double *IPA_SCALE, double *recombination_matrix, double *Q, struct PARAM *p);
void rkck(double *S_sex, double *S_asex, double *IH_sex, double *IH_asex, double *IP_sex, double *IP_asex, double *DSSDT, double *DSADT, double *DIHSDT, double *DIHADT, double *DIPSDT, double *DIPADT, double *SSout, double *SAout, double *IHSout, double *IHAout, double *IPSout, double *IPAout, double *SSerr, double *SAerr, double *IHSerr, double *IHAerr, double *IPSerr, double *IPAerr, double h, double *recombination_matrix, double *Q, struct PARAM *p);
void dynamic(double *SS, double *SA, double *IHS, double *IHA, double *IPS, double *IPA, double *recombination_matrix, double *Q, double *DSSDT, double *DSADT, double *DIHSDT, double *DIHADT, double *DIPSDT, double *DIPADT, struct PARAM *p);
double FMAX(double, double);
double FMIN(double, double);


/*************************************
 * Main function
 *************************************/
void mexFunction(int nlhs, mxArray *plhs[], int nrhs, const mxArray *prhs[])
{
    double *T, *S_sex, *S_asex, *IH_sex, *IH_asex, *IP_sex, *IP_asex, *EQFLAG, *init_pop, *recombination_matrix, *Q, *parameter;
    double *Ttemp, *S_sex_temp, *S_asex_temp, *IH_sex_temp, *IH_asex_temp, *IP_sex_temp, *IP_asex_temp;
    int i, j, k, colLen, maxsteps;
    struct PARAM p;
    
    /* Allocate inputs */
    if(nrhs!=15){
        mexErrMsgTxt("Incorrect number of input arguments!\n");
    }
    else{
        parameter= mxGetPr(prhs[0]);
        p.t_max= *parameter;
        parameter= mxGetPr(prhs[1]);
        p.b= *parameter;
        parameter= mxGetPr(prhs[2]);
        p.c= *parameter;
        parameter= mxGetPr(prhs[3]);
        p.d= *parameter;
        parameter= mxGetPr(prhs[4]);
        p.f= *parameter;   
        parameter= mxGetPr(prhs[5]);        
        p.q= *parameter;
        parameter= mxGetPr(prhs[6]);
        p.alpha= *parameter;
        parameter= mxGetPr(prhs[7]);
        p.beta= *parameter;
        parameter= mxGetPr(prhs[8]);
        p.gamma= *parameter;
        parameter= mxGetPr(prhs[9]);
        p.host_geno_total= (int)*parameter;
        parameter= mxGetPr(prhs[10]);
        p.par_geno_total= (int)*parameter;
        parameter= mxGetPr(prhs[11]);
        p.eqtol= *parameter;
        init_pop= mxGetPr(prhs[12]);
        recombination_matrix= mxGetPr(prhs[13]);
        Q= mxGetPr(prhs[14]);
    }
    maxsteps = (int)MAXSTEPS;
    
    /* Allocate memory */
    Ttemp = malloc(maxsteps*sizeof(double));
    S_sex_temp = malloc(maxsteps*(p.host_geno_total)*sizeof(double));
    S_asex_temp = malloc(maxsteps*(p.host_geno_total)*sizeof(double));
    IH_sex_temp = malloc(maxsteps*(p.host_geno_total)*sizeof(double));
    IH_asex_temp = malloc(maxsteps*(p.host_geno_total)*sizeof(double));
    IP_sex_temp = malloc(maxsteps*(p.par_geno_total)*sizeof(double));
    IP_asex_temp = malloc(maxsteps*(p.par_geno_total)*sizeof(double));
    
    /* Initialise this output */           
    plhs[7] = mxCreateDoubleMatrix(1, 1, mxREAL);
    EQFLAG = mxGetPr(plhs[7]);
    EQFLAG[0] = 0;
    
    /* Call ODE solver */
    colLen = my_rungkut(Ttemp, S_sex_temp, S_asex_temp, IH_sex_temp, IH_asex_temp, IP_sex_temp, IP_asex_temp, EQFLAG, init_pop, recombination_matrix, Q, &p);
    
    /* Create outputs */
    plhs[0] = mxCreateDoubleMatrix(colLen, 1, mxREAL);
    plhs[1] = mxCreateDoubleMatrix(colLen, p.host_geno_total, mxREAL);
    plhs[2] = mxCreateDoubleMatrix(colLen, p.host_geno_total, mxREAL);
    plhs[3] = mxCreateDoubleMatrix(colLen, p.host_geno_total, mxREAL);
    plhs[4] = mxCreateDoubleMatrix(colLen, p.host_geno_total, mxREAL);
    plhs[5] = mxCreateDoubleMatrix(colLen, p.par_geno_total, mxREAL);
    plhs[6] = mxCreateDoubleMatrix(colLen, p.par_geno_total, mxREAL);
    T = mxGetPr(plhs[0]);
    S_sex = mxGetPr(plhs[1]);
    S_asex = mxGetPr(plhs[2]);
    IH_sex = mxGetPr(plhs[3]);
    IH_asex = mxGetPr(plhs[4]);
    IP_sex = mxGetPr(plhs[5]);
    IP_asex = mxGetPr(plhs[6]);
    
    /* Copy data to outputs */
    for (i=0;i<colLen;i++){
        T[i] = Ttemp[i];
        for (j=0;j<p.host_geno_total;j++) {
            S_sex[i + j*colLen] = S_sex_temp[i + j*maxsteps];
            S_asex[i + j*colLen] = S_asex_temp[i + j*maxsteps];
            IH_sex[i + j*colLen] = IH_sex_temp[i + j*maxsteps];
            IH_asex[i + j*colLen] = IH_asex_temp[i + j*maxsteps];
        }
        for (k=0;k<p.par_geno_total;k++) {
            IP_sex[i + k*colLen] = IP_sex_temp[i + k*maxsteps];
            IP_asex[i + k*colLen] = IP_asex_temp[i + k*maxsteps];
        }
    }
    
    /* Free memory */
    free(Ttemp);
    free(S_sex_temp);
    free(S_asex_temp);
    free(IH_sex_temp);
    free(IH_asex_temp);
    free(IP_sex_temp);
    free(IP_asex_temp);
    
    return;
}

/*****************************************
 * ODE solver
 ****************************************/
int my_rungkut (double *T, double *S_sex_out, double *S_asex_out, double *IH_sex_out, double *IH_asex_out, double *IP_sex_out, double *IP_asex_out, double *EQFLAG, double *init_pop, double *recombination_matrix, double *Q, struct PARAM *p){
    
    double *S_sex, *S_asex, *IH_sex, *IH_asex, *IP_sex, *IP_asex, *DSSDT, *DSADT, *DIHSDT, *DIHADT, *DIPSDT, *DIPADT, *SS_SCALE, *SA_SCALE, *IHS_SCALE, *IHA_SCALE, *IPS_SCALE, *IPA_SCALE;
    double *SSMIN, *SSMAX, *SAMIN, *SAMAX, *IHSMIN, *IHSMAX, *IHAMIN, *IHAMAX, *IPSMIN, *IPSMAX, *IPAMIN, *IPAMAX, hnext[1], h[1];
    double t, nextcheck;
    int i, j, k, exitflag, count, maxsteps;
    
    /* Allocate memory */
    S_sex = malloc(p->host_geno_total*sizeof(double));
    S_asex = malloc(p->host_geno_total*sizeof(double));
    IH_sex = malloc(p->host_geno_total*sizeof(double));
    IH_asex = malloc(p->host_geno_total*sizeof(double));
    IP_sex = malloc(p->par_geno_total*sizeof(double));
    IP_asex = malloc(p->par_geno_total*sizeof(double));    
    DSSDT = malloc(p->host_geno_total*sizeof(double));
    DSADT = malloc(p->host_geno_total*sizeof(double));
    DIHSDT = malloc(p->host_geno_total*sizeof(double));
    DIHADT = malloc(p->host_geno_total*sizeof(double));
    DIPSDT = malloc(p->par_geno_total*sizeof(double));
    DIPADT = malloc(p->par_geno_total*sizeof(double));    
    SS_SCALE = malloc(p->host_geno_total*sizeof(double));
    SA_SCALE = malloc(p->host_geno_total*sizeof(double));
    IHS_SCALE = malloc(p->host_geno_total*sizeof(double));
    IHA_SCALE = malloc(p->host_geno_total*sizeof(double));
    IPS_SCALE = malloc(p->par_geno_total*sizeof(double));
    IPA_SCALE = malloc(p->par_geno_total*sizeof(double));    
    SSMIN = malloc(p->host_geno_total*sizeof(double));
    SAMIN = malloc(p->host_geno_total*sizeof(double));
    IHSMIN = malloc(p->host_geno_total*sizeof(double));
    IHAMIN = malloc(p->host_geno_total*sizeof(double));
    IPSMIN = malloc(p->par_geno_total*sizeof(double));
    IPAMIN = malloc(p->par_geno_total*sizeof(double));    
    SSMAX = malloc(p->host_geno_total*sizeof(double));
    SAMAX = malloc(p->host_geno_total*sizeof(double));
    IHSMAX = malloc(p->host_geno_total*sizeof(double));
    IHAMAX = malloc(p->host_geno_total*sizeof(double));
    IPSMAX = malloc(p->par_geno_total*sizeof(double));
    IPAMAX = malloc(p->par_geno_total*sizeof(double));
    
    /* Other parameters */
    exitflag = 1;
    count=0;
    k=1;
    h[0] = 1e-3;
    hnext[0] = 1e-3;
    t=0;
    nextcheck = INTERVAL;
    maxsteps = (int)MAXSTEPS;
    
    /* Initialise populations */
    for(i=0;i<p->host_geno_total;i++){
        S_sex[i] = init_pop[i];
        S_asex[i] = init_pop[p->host_geno_total+i];
        IH_sex[i] = init_pop[2*p->host_geno_total+i];
        IH_asex[i] = init_pop[3*p->host_geno_total+i];
    }
    for(j=0;j<p->par_geno_total;j++){
        IP_sex[j] = init_pop[4*p->host_geno_total + j];
        IP_asex[j] = init_pop[4*p->host_geno_total + p->par_geno_total + j];
    }
    
    /* Initialise equilibrium arrays */
    for(i=0;i<p->host_geno_total;i++){
        SSMIN[i] = S_sex[i];
        SSMAX[i] = S_sex[i];
        SAMIN[i] = S_asex[i];
        SAMAX[i] = S_asex[i];
        IHSMIN[i] = IH_sex[i];
        IHSMAX[i] = IH_sex[i];
        IHAMIN[i] = IH_asex[i];
        IHAMAX[i] = IH_asex[i];
    }
    for(j=0;j<p->par_geno_total;j++){
        IPSMIN[j] = IP_sex[j];
        IPSMAX[j] = IP_sex[j];
        IPAMIN[j] = IP_asex[j];
        IPAMAX[j] = IP_asex[j];
    }
    
    /* Update output */
    T[0]=t;
    for (i=0; i<p->host_geno_total; i++) {
        S_sex_out[i*maxsteps] = S_sex[i];
        S_asex_out[i*maxsteps] = S_asex[i];
        IH_sex_out[i*maxsteps] = IH_sex[i];
        IH_asex_out[i*maxsteps] = IH_asex[i];
    }
    for(j=0;j<p->par_geno_total;j++){
        IP_sex_out[j*maxsteps] = IP_sex[j];
        IP_asex_out[j*maxsteps] = IP_asex[j];
    }
    
    /* Main loop: */
    do{
        /* This ensures the final step lands us on the final time point */
        if(1.1*hnext[0]>(p->t_max-t)){
            hnext[0] = p->t_max-t;
            h[0] = p->t_max-t;
            t=p->t_max;
            exitflag=0;
        }
        else{
            h[0] = hnext[0];
            t+=h[0];
        }
        if(t>=p->t_max) {
            t=p->t_max;
            exitflag=0;
        }
        /* This is where the equations are first solved */
        dynamic(S_sex, S_asex, IH_sex, IH_asex, IP_sex, IP_asex, recombination_matrix, Q, DSSDT, DSADT, DIHSDT, DIHADT, DIPSDT, DIPADT, p);

        /* Adjust the step size to maintain accuracy */
        for (i=0; i<p->host_geno_total; i++){
            S_sex[i] = FMAX(S_sex[i],0);
            S_asex[i] = FMAX(S_asex[i],0);
            IH_sex[i] = FMAX(IH_sex[i],0);
            IH_asex[i] = FMAX(IH_asex[i],0);
            SS_SCALE[i]=fabs(S_sex[i])+fabs(DSSDT[i]*(*h))+TINY;
            SA_SCALE[i]=fabs(S_asex[i])+fabs(DSADT[i]*(*h))+TINY;
            IHS_SCALE[i]=fabs(IH_sex[i])+fabs(DIHSDT[i]*(*h))+TINY;
            IHA_SCALE[i]=fabs(IH_asex[i])+fabs(DIHADT[i]*(*h))+TINY;
        }
        for(j=0;j<p->par_geno_total;j++){
            IP_sex[j] = FMAX(IP_sex[j],0);
            IP_asex[j] = FMAX(IP_asex[j],0);
            IPS_SCALE[j]=fabs(IP_sex[j])+fabs(DIPSDT[j]*(*h))+TINY;
            IPA_SCALE[j]=fabs(IP_asex[j])+fabs(DIPADT[j]*(*h))+TINY;
        }
        
        /* RK solver & adaptive step-size */
        rkqs(S_sex, S_asex, IH_sex, IH_asex, IP_sex, IP_asex, DSSDT, DSADT, DIHSDT, DIHADT, DIPSDT, DIPADT, h, hnext, SS_SCALE, SA_SCALE, IHS_SCALE, IHA_SCALE, IPS_SCALE, IPA_SCALE, recombination_matrix, Q, p);
        
        /* Make sure nothin has gone negative */
        for (i=0; i<p->host_geno_total; i++){
            S_sex[i] = FMAX(S_sex[i],0);
            S_asex[i] = FMAX(S_asex[i],0);
            IH_sex[i] = FMAX(IH_sex[i],0);
            IH_asex[i] = FMAX(IH_asex[i],0);
        }
        for(j=0;j<p->par_geno_total;j++){
            IP_sex[j] = FMAX(IP_sex[j],0);
            IP_asex[j] = FMAX(IP_asex[j],0);
        }
        
        /* Update output */
        count++;
        T[count] = t;
        for (i=0; i<p->host_geno_total; i++) {
            S_sex_out[count + i*maxsteps] = S_sex[i];
            S_asex_out[count + i*maxsteps] = S_asex[i];
            IH_sex_out[count + i*maxsteps] = IH_sex[i];
            IH_asex_out[count + i*maxsteps] = IH_asex[i];
        }
        for(j=0;j<p->par_geno_total;j++){
            IP_sex_out[count + j*maxsteps] = IP_sex[j];
            IP_asex_out[count + j*maxsteps] = IP_asex[j];
        }
    
        /* For equilibrium check */
        for (i=0; i<p->host_geno_total; i++){
            SSMIN[i] = FMIN(SSMIN[i],S_sex[i]);
            SAMIN[i] = FMIN(SAMIN[i],S_asex[i]);
            SSMAX[i] = FMAX(SSMAX[i],S_sex[i]);
            SAMAX[i] = FMAX(SAMAX[i],S_asex[i]);
            IHSMIN[i] = FMIN(IHSMIN[i],IH_sex[i]);
            IHAMIN[i] = FMIN(IHAMIN[i],IH_asex[i]);
            IHSMAX[i] = FMAX(IHSMAX[i],IH_sex[i]);
            IHAMAX[i] = FMAX(IHAMAX[i],IH_asex[i]);
        }
        for(j=0;j<p->par_geno_total;j++){
            IPSMIN[j] = FMIN(IPSMIN[j],IP_sex[j]);
            IPAMIN[j] = FMIN(IPAMIN[j],IP_asex[j]);
            IPSMAX[j] = FMAX(IPSMAX[j],IP_sex[j]);
            IPAMAX[j] = FMAX(IPAMAX[j],IP_asex[j]);
        }
        
        /* Check if we're close to equilibrium */
        if(t>nextcheck){
            exitflag = 0;
            for (i=0; i<p->host_geno_total; i++){
                if(fabs(SSMAX[i]-SSMIN[i])>p->eqtol || fabs(SAMAX[i]-SAMIN[i])>p->eqtol || fabs(IHSMAX[i]-IHSMIN[i])>p->eqtol || fabs(IHAMAX[i]-IHAMIN[i])>p->eqtol){
                    exitflag = 1;
                    break;
                }
            }
            if(exitflag==0){
                for(j=0;j<p->par_geno_total;j++){
                    if(fabs(IPSMAX[j]-IPSMIN[j])>p->eqtol || fabs(IPAMAX[j]-IPAMIN[j])>p->eqtol){
                        exitflag = 1;
                        break;
                    }
                }
            }
            /* If close to equilibrium, then break */
            if(exitflag==0){
                t=p->t_max;
                T[count] = t;
                EQFLAG[0] = 1;
                break;
            }
            
            /* If not, then reset min/max values for each class */
            nextcheck+=INTERVAL;
            for (i=0; i<p->host_geno_total; i++){
                SSMIN[i] = S_sex[i];
                SAMIN[i] = S_asex[i];
                SSMAX[i] = S_sex[i];
                SAMAX[i] = S_asex[i];
                IHSMIN[i] = IH_sex[i];
                IHAMIN[i] = IH_asex[i];
                IHSMAX[i] = IH_sex[i];
                IHAMAX[i] = IH_asex[i];
            }
            for(j=0;j<p->par_geno_total;j++){
                IPSMIN[j] = IP_sex[j];
                IPAMIN[j] = IP_asex[j];
                IPSMAX[j] = IP_sex[j];
                IPAMAX[j] = IP_asex[j];
            }
        }
    }while(count<(maxsteps-1) && t<=p->t_max && exitflag);
    count++;
    
    /* Free memory */
    free(S_sex);
    free(S_asex);
    free(IH_sex);
    free(IH_asex);
    free(IP_sex);
    free(IP_asex);
    free(DSSDT);
    free(DSADT);
    free(DIHSDT);
    free(DIHADT);
    free(DIPSDT);
    free(DIPADT);
    free(SS_SCALE);
    free(SA_SCALE);
    free(IHS_SCALE);
    free(IHA_SCALE);
    free(IPS_SCALE);
    free(IPA_SCALE);
    free(SSMIN);
    free(SAMIN);
    free(IHSMIN);
    free(IHAMIN);
    free(IPSMIN);
    free(IPAMIN);
    free(SSMAX);
    free(SAMAX);
    free(IHSMAX);
    free(IHAMAX);
    free(IPSMAX);
    free(IPAMAX);
    
    return count;
}

/***************************************
 * This generates the adaptive step-size
 **************************************/
void rkqs(double *S_sex, double *S_asex, double *IH_sex, double *IH_asex, double *IP_sex, double *IP_asex, double *DSSDT, double *DSADT, double *DIHSDT, double *DIHADT, double *DIPSDT, double *DIPADT, double *h, double *hnext, double *SS_SCALE, double *SA_SCALE, double *IHS_SCALE, double *IHA_SCALE, double *IPS_SCALE, double *IPA_SCALE, double *recombination_matrix, double *Q, struct PARAM *p)
{
    double *S_sex_temp, *S_asex_temp, *IH_sex_temp, *IH_asex_temp, *IP_sex_temp, *IP_asex_temp, *S_sex_err, *S_asex_err, *IH_sex_err, *IH_asex_err, *IP_sex_err, *IP_asex_err;
    double htemp, errmax;
    int i, j, count;
    
    /* Allocate memory */
    S_sex_temp = malloc(p->host_geno_total*sizeof(double));
    S_asex_temp = malloc(p->host_geno_total*sizeof(double));
    IH_sex_temp = malloc(p->host_geno_total*sizeof(double));
    IH_asex_temp = malloc(p->host_geno_total*sizeof(double));
    IP_sex_temp = malloc(p->par_geno_total*sizeof(double));
    IP_asex_temp = malloc(p->par_geno_total*sizeof(double));
    S_sex_err = malloc(p->host_geno_total*sizeof(double));
    S_asex_err = malloc(p->host_geno_total*sizeof(double));
    IH_sex_err = malloc(p->host_geno_total*sizeof(double));
    IH_asex_err = malloc(p->host_geno_total*sizeof(double));
    IP_sex_err = malloc(p->par_geno_total*sizeof(double));
    IP_asex_err = malloc(p->par_geno_total*sizeof(double));
    
    count = 0;
    for(;;)
    {
        rkck(S_sex, S_asex, IH_sex, IH_asex, IP_sex, IP_asex, DSSDT, DSADT, DIHSDT, DIHADT, DIPSDT, DIPADT, S_sex_temp, S_asex_temp, IH_sex_temp, IH_asex_temp, IP_sex_temp, IP_asex_temp, S_sex_err, S_asex_err, IH_sex_err, IH_asex_err, IP_sex_err, IP_asex_err, *h, recombination_matrix, Q, p);
        
        errmax= 0.0;
        for(i=0;i<p->host_geno_total;i++){
            errmax= FMAX(errmax, fabs(S_sex_err[i]/(SS_SCALE[i])));
            errmax= FMAX(errmax, fabs(S_asex_err[i]/(SA_SCALE[i]))); 
            errmax= FMAX(errmax, fabs(IH_sex_err[i]/(IHS_SCALE[i])));
            errmax= FMAX(errmax, fabs(IH_asex_err[i]/(IHA_SCALE[i])));   
        }
        for(j=0;j<p->par_geno_total;j++){
            errmax= FMAX(errmax, fabs(IP_sex_err[j]/(IPS_SCALE[j])));
            errmax= FMAX(errmax, fabs(IP_asex_err[j]/(IPA_SCALE[j])));
        }
        errmax/= EPS;
        if(errmax<=1.0) break;
        htemp= 0.9*(*h)*pow(errmax, -0.25);
        *h= (*h>=0.0 ? FMAX(htemp, 0.1*(*h)) : FMIN(htemp, 0.1*(*h)));
        count++;
            
        if(count>1e4){
            printf("%f\n",errmax);
            mexErrMsgTxt("stuck in loop!\n");
            break;
        }
    }    
    if(errmax > 1.89E-4) {
        *hnext= 0.9*(*h)*pow(errmax, -0.2);
    }
    else {
        *hnext= 5.0*(*h);
    }    
    *hnext = FMAX(*hnext, p->t_max/MAXSTEPS);
    
    for(i=0;i<p->host_geno_total;i++){
        S_sex[i] = S_sex_temp[i];
        S_asex[i] = S_asex_temp[i];
        IH_sex[i] = IH_sex_temp[i];
        IH_asex[i] = IH_asex_temp[i];
    }
    for(j=0;j<p->par_geno_total;j++){
        IP_sex[j] = IP_sex_temp[j];
        IP_asex[j] = IP_asex_temp[j];
    }
    
    /* Free memory */
    free(S_sex_temp);
    free(S_asex_temp);
    free(IH_sex_temp);
    free(IH_asex_temp);
    free(IP_sex_temp);
    free(IP_asex_temp);
    free(S_sex_err);
    free(S_asex_err);
    free(IH_sex_err);
    free(IH_asex_err);
    free(IP_sex_err);
    free(IP_asex_err);
}

/**************************************
 * Standard RK solver
 **************************************/
void rkck(double *S_sex, double *S_asex, double *IH_sex, double *IH_asex, double *IP_sex, double *IP_asex, double *DSSDT, double *DSADT, double *DIHSDT, double *DIHADT, double *DIPSDT, double *DIPADT, double *SSout, double *SAout, double *IHSout, double *IHAout, double *IPSout, double *IPAout, double *SSerr, double *SAerr, double *IHSerr, double *IHAerr, double *IPSerr, double *IPAerr, double h, double *recombination_matrix, double *Q, struct PARAM *p){
    int i, j;
    double *SSk1, *SSk2, *SSk3, *SSk4, *SSk5, *SSk6, *SStemp;
    double *SAk1, *SAk2, *SAk3, *SAk4, *SAk5, *SAk6, *SAtemp;
    double *IHSk1, *IHSk2, *IHSk3, *IHSk4, *IHSk5, *IHSk6, *IHStemp;
    double *IHAk1, *IHAk2, *IHAk3, *IHAk4, *IHAk5, *IHAk6, *IHAtemp;
    double *IPSk1, *IPSk2, *IPSk3, *IPSk4, *IPSk5, *IPSk6, *IPStemp;
    double *IPAk1, *IPAk2, *IPAk3, *IPAk4, *IPAk5, *IPAk6, *IPAtemp;
    double dc1=c1-2825.0/27648.0, dc3=c3-18575.0/48384.0, dc4=c4-13525.0/55296.0, 
            dc6=c6-0.25;
    
    /* Allocate memory */    
    SSk1 = malloc(p->host_geno_total*sizeof(double));
    SSk2 = malloc(p->host_geno_total*sizeof(double));
    SSk3 = malloc(p->host_geno_total*sizeof(double));
    SSk4 = malloc(p->host_geno_total*sizeof(double));
    SSk5 = malloc(p->host_geno_total*sizeof(double));
    SSk6 = malloc(p->host_geno_total*sizeof(double));
    SStemp = malloc(p->host_geno_total*sizeof(double));
    
    SAk1 = malloc(p->host_geno_total*sizeof(double));
    SAk2 = malloc(p->host_geno_total*sizeof(double));
    SAk3 = malloc(p->host_geno_total*sizeof(double));
    SAk4 = malloc(p->host_geno_total*sizeof(double));
    SAk5 = malloc(p->host_geno_total*sizeof(double));
    SAk6 = malloc(p->host_geno_total*sizeof(double));
    SAtemp = malloc(p->host_geno_total*sizeof(double));
    
    IHSk1 = malloc(p->host_geno_total*sizeof(double));
    IHSk2 = malloc(p->host_geno_total*sizeof(double));
    IHSk3 = malloc(p->host_geno_total*sizeof(double));
    IHSk4 = malloc(p->host_geno_total*sizeof(double));
    IHSk5 = malloc(p->host_geno_total*sizeof(double));
    IHSk6 = malloc(p->host_geno_total*sizeof(double));
    IHStemp = malloc(p->host_geno_total*sizeof(double));
    
    IHAk1 = malloc(p->host_geno_total*sizeof(double));
    IHAk2 = malloc(p->host_geno_total*sizeof(double));
    IHAk3 = malloc(p->host_geno_total*sizeof(double));
    IHAk4 = malloc(p->host_geno_total*sizeof(double));
    IHAk5 = malloc(p->host_geno_total*sizeof(double));
    IHAk6 = malloc(p->host_geno_total*sizeof(double));  
    IHAtemp = malloc(p->host_geno_total*sizeof(double));  
    
    IPSk1 = malloc(p->par_geno_total*sizeof(double));
    IPSk2 = malloc(p->par_geno_total*sizeof(double));
    IPSk3 = malloc(p->par_geno_total*sizeof(double));
    IPSk4 = malloc(p->par_geno_total*sizeof(double));
    IPSk5 = malloc(p->par_geno_total*sizeof(double));
    IPSk6 = malloc(p->par_geno_total*sizeof(double));
    IPStemp = malloc(p->par_geno_total*sizeof(double));
    
    IPAk1 = malloc(p->par_geno_total*sizeof(double));
    IPAk2 = malloc(p->par_geno_total*sizeof(double));
    IPAk3 = malloc(p->par_geno_total*sizeof(double));
    IPAk4 = malloc(p->par_geno_total*sizeof(double));
    IPAk5 = malloc(p->par_geno_total*sizeof(double));
    IPAk6 = malloc(p->par_geno_total*sizeof(double));  
    IPAtemp = malloc(p->par_geno_total*sizeof(double));  
    
    for(i=0;i<p->host_geno_total;i++){
        SStemp[i] = S_sex[i] + b21*h*DSSDT[i];
        SAtemp[i] = S_asex[i] + b21*h*DSADT[i];
        IHStemp[i] = IH_sex[i] + b21*h*DIHSDT[i];
        IHAtemp[i] = IH_asex[i] + b21*h*DIHADT[i];
    }
    for(j=0;j<p->par_geno_total;j++){
        IPStemp[j] = IP_sex[j] + b21*h*DIPSDT[j];
        IPAtemp[j] = IP_asex[j] + b21*h*DIPADT[j];
    }
    dynamic(SStemp, SAtemp, IHStemp, IHAtemp,  IPStemp, IPAtemp, recombination_matrix, Q, SSk2, SAk2, IHSk2, IHAk2, IPSk2, IPAk2, p);
    
    for(i=0;i<p->host_geno_total;i++){
        SStemp[i] = S_sex[i]+h*(b31*DSSDT[i]+b32*SSk2[i]);
        SAtemp[i] = S_asex[i]+h*(b31*DSADT[i]+b32*SAk2[i]);
        IHStemp[i] = IH_sex[i]+h*(b31*DIHSDT[i]+b32*IHSk2[i]);
        IHAtemp[i] = IH_asex[i]+h*(b31*DIHADT[i]+b32*IHAk2[i]);
    }    
    for(j=0;j<p->par_geno_total;j++){
        IPStemp[j] = IP_sex[j]+h*(b31*DIPSDT[j]+b32*IPSk2[j]);
        IPAtemp[j] = IP_asex[j]+h*(b31*DIPADT[j]+b32*IPAk2[j]);
    }
    dynamic(SStemp, SAtemp, IHStemp, IHAtemp,  IPStemp, IPAtemp, recombination_matrix, Q, SSk3, SAk3, IHSk3, IHAk3, IPSk3, IPAk3, p);
    
    for(i=0;i<p->host_geno_total;i++){
        SStemp[i] = S_sex[i]+h*(b41*DSSDT[i]+b42*SSk2[i]+b43*SSk3[i]);
        SAtemp[i] = S_asex[i]+h*(b41*DSADT[i]+b42*SAk2[i]+b43*SAk3[i]);
        IHStemp[i] = IH_sex[i]+h*(b41*DIHSDT[i]+b42*IHSk2[i]+b43*IHSk3[i]);
        IHAtemp[i] = IH_asex[i]+h*(b41*DIHADT[i]+b42*IHAk2[i]+b43*IHAk3[i]);
    }
    for(j=0;j<p->par_geno_total;j++){
        IPStemp[j] = IP_sex[j]+h*(b41*DIPSDT[j]+b42*IPSk2[j]+b43*IPSk3[j]);
        IPAtemp[j] = IP_asex[j]+h*(b41*DIPADT[j]+b42*IPAk2[j]+b43*IPAk3[j]);
    }
    dynamic(SStemp, SAtemp, IHStemp, IHAtemp,  IPStemp, IPAtemp, recombination_matrix, Q, SSk4, SAk4, IHSk4, IHAk4, IPSk4, IPAk4, p);
    
    for(i=0;i<p->host_geno_total;i++){
        SStemp[i] = S_sex[i]+h*(b51*DSSDT[i]+b52*SSk2[i]+b53*SSk3[i]+b54*SSk4[i]);
        SAtemp[i] = S_asex[i]+h*(b51*DSADT[i]+b52*SAk2[i]+b53*SAk3[i]+b54*SAk4[i]);
        IHStemp[i] = IH_sex[i]+h*(b51*DIHSDT[i]+b52*IHSk2[i]+b53*IHSk3[i]+b54*IHSk4[i]);
        IHAtemp[i] = IH_asex[i]+h*(b51*DIHADT[i]+b52*IHAk2[i]+b53*IHAk3[i]+b54*IHAk4[i]);
    }
    for(j=0;j<p->par_geno_total;j++){
        IPStemp[j] = IP_sex[j]+h*(b51*DIPSDT[j]+b52*IPSk2[j]+b53*IPSk3[j]+b54*IPSk4[j]);
        IPAtemp[j] = IP_asex[j]+h*(b51*DIPADT[j]+b52*IPAk2[j]+b53*IPAk3[j]+b54*IPAk4[j]);
    }
    dynamic(SStemp, SAtemp, IHStemp, IHAtemp,  IPStemp, IPAtemp, recombination_matrix, Q, SSk5, SAk5, IHSk5, IHAk5, IPSk5, IPAk5, p);
    
    for(i=0;i<p->host_geno_total;i++){
        SStemp[i] = S_sex[i]+h*(b61*DSSDT[i]+b62*SSk2[i]+b63*SSk3[i]+b64*SSk4[i]+b65*SSk5[i]);
        SAtemp[i] = S_asex[i]+h*(b61*DSADT[i]+b62*SAk2[i]+b63*SAk3[i]+b64*SAk4[i]+b65*SAk5[i]);        
        IHStemp[i] = IH_sex[i]+h*(b61*DIHSDT[i]+b62*IHSk2[i]+b63*IHSk3[i]+b64*IHSk4[i]+b65*IHSk5[i]);
        IHAtemp[i] = IH_asex[i]+h*(b61*DIHADT[i]+b62*IHAk2[i]+b63*IHAk3[i]+b64*IHAk4[i]+b65*IHAk5[i]);
    }
    for(j=0;j<p->par_geno_total;j++){
        IPStemp[j] = IP_sex[j]+h*(b61*DIPSDT[j]+b62*IPSk2[j]+b63*IPSk3[j]+b64*IPSk4[j]+b65*IPSk5[j]);
        IPAtemp[j] = IP_asex[j]+h*(b61*DIPADT[j]+b62*IPAk2[j]+b63*IPAk3[j]+b64*IPAk4[j]+b65*IPAk5[j]);
    }
    dynamic(SStemp, SAtemp, IHStemp, IHAtemp,  IPStemp, IPAtemp, recombination_matrix, Q, SSk6, SAk6, IHSk6, IHAk6, IPSk6, IPAk6, p);
    
    for(i=0;i<p->host_geno_total;i++){
        SSout[i]= S_sex[i]+h*(c1*DSSDT[i]+c3*SSk3[i]+c4*SSk4[i]+c6*SSk6[i]);
        SSerr[i]= h*(dc1*DSSDT[i]+dc3*SSk3[i]+dc4*SSk4[i]+dc5*SSk5[i]+dc6*SSk6[i]);
        SAout[i]= S_asex[i]+h*(c1*DSADT[i]+c3*SAk3[i]+c4*SAk4[i]+c6*SAk6[i]);
        SAerr[i]= h*(dc1*DSADT[i]+dc3*SAk3[i]+dc4*SAk4[i]+dc5*SAk5[i]+dc6*SAk6[i]);
        IHSout[i]= IH_sex[i]+h*(c1*DIHSDT[i]+c3*IHSk3[i]+c4*IHSk4[i]+c6*IHSk6[i]);
        IHSerr[i]= h*(dc1*DIHSDT[i]+dc3*IHSk3[i]+dc4*IHSk4[i]+dc5*IHSk5[i]+dc6*IHSk6[i]);
        IHAout[i]= IH_asex[i]+h*(c1*DIHADT[i]+c3*IHAk3[i]+c4*IHAk4[i]+c6*IHAk6[i]);
        IHAerr[i]= h*(dc1*DIHADT[i]+dc3*IHAk3[i]+dc4*IHAk4[i]+dc5*IHAk5[i]+dc6*IHAk6[i]);
    }
    for(j=0;j<p->par_geno_total;j++){
        IPSout[j]= IP_sex[j]+h*(c1*DIPSDT[j]+c3*IPSk3[j]+c4*IPSk4[j]+c6*IPSk6[j]);
        IPSerr[j]= h*(dc1*DIPSDT[j]+dc3*IPSk3[j]+dc4*IPSk4[j]+dc5*IPSk5[j]+dc6*IPSk6[j]);
        IPAout[j]= IP_asex[j]+h*(c1*DIPADT[j]+c3*IPAk3[j]+c4*IPAk4[j]+c6*IPAk6[j]);
        IPAerr[j]= h*(dc1*DIPADT[j]+dc3*IPAk3[j]+dc4*IPAk4[j]+dc5*IPAk5[j]+dc6*IPAk6[j]);
    }

    /* Free memory */
    free(SSk1);
    free(SSk2);
    free(SSk3);
    free(SSk4);
    free(SSk5);
    free(SSk6);
    free(SAk1);
    free(SAk2);
    free(SAk3);
    free(SAk4);
    free(SAk5);
    free(SAk6);
    free(SStemp);
    free(SAtemp);
    free(IHSk1);
    free(IHSk2);
    free(IHSk3);
    free(IHSk4);
    free(IHSk5);
    free(IHSk6);
    free(IHAk1);
    free(IHAk2);
    free(IHAk3);
    free(IHAk4);
    free(IHAk5);
    free(IHAk6);
    free(IHStemp);
    free(IHAtemp);
    free(IPSk1);
    free(IPSk2);
    free(IPSk3);
    free(IPSk4);
    free(IPSk5);
    free(IPSk6);
    free(IPAk1);
    free(IPAk2);
    free(IPAk3);
    free(IPAk4);
    free(IPAk5);
    free(IPAk6);
    free(IPStemp);
    free(IPAtemp);
}

/**************************************
 * Population and evolutionary dynamics
 **************************************/
void dynamic(double *SS, double *SA, double *IHS, double *IHA, double *IPS, double *IPA, double *recombination_matrix, double *Q, double *DSSDT, double *DSADT, double *DIHSDT, double *DIHADT, double *DIPSDT, double *DIPADT, struct PARAM *p){
    
    int i, j;
    double hostsum_sex, hostsum_asex, hostsum, Fsum;
    double *gametes, *zygotes, *F, *births_sex, *births_asex, *infections_sex, *infections_asex, *infections_sex_H, *infections_asex_H, *infections_sex_P, *infections_asex_P;
    
    /* Allocate memory */
    F = malloc(p->host_geno_total*sizeof(double));
    gametes = malloc(p->par_geno_total*sizeof(double));
    zygotes = malloc(p->host_geno_total*sizeof(double));
    births_sex = malloc(p->host_geno_total*sizeof(double));
    births_asex = malloc(p->host_geno_total*sizeof(double));
    infections_sex = malloc(p->host_geno_total*p->par_geno_total*sizeof(double));
    infections_asex = malloc(p->host_geno_total*p->par_geno_total*sizeof(double));
    infections_sex_H = malloc(p->host_geno_total*sizeof(double));
    infections_asex_H = malloc(p->host_geno_total*sizeof(double));
    infections_sex_P = malloc(p->par_geno_total*sizeof(double));
    infections_asex_P = malloc(p->par_geno_total*sizeof(double));
    
    /* Population sums */
    hostsum_sex = 0;
    hostsum_asex = 0;
    Fsum = 0;
    for(i=0;i<p->host_geno_total;i++){
        hostsum_sex += SS[i] + IHS[i];
        hostsum_asex += SA[i] + IHA[i];
        F[i] = SS[i] + p->f*IHS[i];
        Fsum += F[i];
    }    
    hostsum = hostsum_sex + hostsum_asex;
    
    /* Sexual births */
    /* Pool of gametes */
    for(j=0;j<p->par_geno_total;j++){ // This is also equal to the number of host haplotypes
        gametes[j] = 0;
        for(i=0;i<p->host_geno_total;i++){
            gametes[j] += FMIN(F[i]/FMAX(Fsum,1E-30),1)*recombination_matrix[j*p->host_geno_total + i];
        }
    }
    
    /* Create zygotes based on pool of gametes */
    for(i=0;i<p->par_geno_total;i++){
        for(j=0;j<p->par_geno_total;j++){
            zygotes[j + i*p->par_geno_total] = gametes[i]*gametes[j];
        }
    }
    for(i=0;i<p->host_geno_total;i++){
        births_sex[i] = FMAX(0,p->b*p->c*(1-p->q*hostsum)*zygotes[i]*Fsum);
    }
    
    /* Asex births */    
    for(i=0;i<p->host_geno_total;i++){
        births_asex[i] = FMAX(0,p->b*(1-p->q*hostsum)*(SA[i] + p->f*IHA[i]));
    }
    
    /* Infections */
    for(j=0;j<p->par_geno_total;j++){
        infections_sex_P[j] = 0;
        infections_asex_P[j] = 0;
    }
    for(i=0;i<p->host_geno_total;i++){
        infections_sex_H[i] = 0;
        infections_asex_H[i] = 0;
        for(j=0;j<p->par_geno_total;j++){
            infections_sex[i + j*p->host_geno_total] = p->beta*Q[i + j*p->host_geno_total]*SS[i]*(IPS[j]+IPA[j]+TINY2);
            infections_asex[i + j*p->host_geno_total] = p->beta*Q[i + j*p->host_geno_total]*SA[i]*(IPS[j]+IPA[j]+TINY2);
            infections_sex_H[i] += infections_sex[i + j*p->host_geno_total];
            infections_asex_H[i] += infections_asex[i + j*p->host_geno_total];
            infections_sex_P[j] += infections_sex[i + j*p->host_geno_total];
            infections_asex_P[j] += infections_asex[i + j*p->host_geno_total];            
        }
    }
    
    /* ODEs */
    for(i=0;i<p->host_geno_total;i++){
        DSSDT[i] = births_sex[i] - infections_sex_H[i] - p->d*SS[i] + p->gamma*IHS[i];
        DSADT[i] = births_asex[i] - infections_asex_H[i] - p->d*SA[i] + p->gamma*IHA[i];
        DIHSDT[i] = infections_sex_H[i] - (p->d + p->alpha + p->gamma)*IHS[i];
        DIHADT[i] = infections_asex_H[i] - (p->d + p->alpha + p->gamma)*IHA[i];
    }
    for(j=0;j<p->par_geno_total;j++){
        DIPSDT[j] = infections_sex_P[j] - (p->d + p->alpha + p->gamma)*IPS[j];
        DIPADT[j] = infections_asex_P[j] - (p->d + p->alpha + p->gamma)*IPA[j];
    }
    
    /* Free memory */
    free(F);
    free(gametes);
    free(zygotes);
    free(births_sex);
    free(births_asex);
    free(infections_sex);    
    free(infections_asex);    
    free(infections_sex_H);    
    free(infections_asex_H); 
    free(infections_sex_P);    
    free(infections_asex_P);
}

/***************************************
 * Return maximum of two inputs
 ***************************************/
double FMAX(double l, double r)
{
    if(l>r)return l;
    else   return r;
}

/***************************************
 * Return minimum of two inputs
 ***************************************/
double FMIN(double l, double r)
{
    if(l<r)return l;
    else   return r;
}
```
